# Machine Learning Reveals Lipidome Remodeling Dynamics in a Mouse Model of Ovarian Cancer

**DOI:** 10.1101/2023.01.04.520434

**Authors:** Olatomiwa O. Bifarin, Samyukta Sah, David A. Gaul, Samuel G. Moore, Ruihong Chen, Murugesan Palaniappan, Jaeyeon Kim, Martin M. Matzuk, Facundo M. Fernández

## Abstract

Ovarian cancer (OC) is one of the deadliest cancers affecting the female reproductive system. It may present little or no symptoms at the early stages, and typically unspecific symptoms at later stages. High-grade serous ovarian cancer (HGSC) is the subtype responsible for most ovarian cancer deaths. However, very little is known about the metabolic course of this disease, particularly in its early stages. In this longitudinal study, we examined the temporal course of serum lipidome changes using a robust HGSC mouse model and machine learning data analysis. Early progression of HGSC was marked by increased levels of phosphatidylcholines and phosphatidylethanolamines. In contrast, later stages featured more diverse lipids alterations, including fatty acids and their derivatives, triglycerides, ceramides, hexosylceramides, sphingomyelins, lysophosphatidylcholines, and phosphatidylinositols. These alterations underscored unique perturbations in cell membrane stability, proliferation, and survival during cancer development and progression, offering potential targets for early detection and prognosis of human ovarian cancer.

**Teaser:** Time-resolved lipidome remodeling in an ovarian cancer model is studied through lipidomics and machine learning.

## Introduction

The absence of reliable non-invasive ovarian cancer (OC) diagnostics leads to more deaths than any other cancer associated with the female reproductive system, with 419,085 deaths from 1990 to 2019 in the United States alone (*1*). It is the fifth leading cause of cancer-related death in women (*2*). Failure of early detection remains the most daunting challenge in OC diagnosis (*3*). In the United States, the 5-year survival rate is 93.1% for localized OC, but it is reduced drastically to only 30.8% for metastatic OC (*4*). High-grade serous ovarian cancer (HGSC) is the most frequent subtype accounting for 70-80% of all OC deaths (*5, 6*). Early diagnosis is therefore imperative for reducing OC mortality. However, OC often eludes detection until an advanced stage (*6*) and the molecular pathogenesis underlying early-stage OC remains poorly understood. To study the biochemical underpinnings of early-stage OC pathogenesis, we conducted in-depth lipidomic analyses in a *Dicer1-Pten* double-knockout (DKO) mouse model as a function of time. These mice faithfully recapitulate human HGSC with phenotypic, histopathologic, and molecular similarities (*7, 8*) and exhibit stepwise development and progression of HGSC, beginning with a premalignant phase, tumor initiation, and malignant growth in the primary tissue before advancing to early metastases, widespread metastases, and ultimately death.

It is now widely accepted that cancer is a metabolic disease (*9*). As such, metabolomics/lipidomics are central to cancer biology. Metabolomics and lipidomics allow for measuring and identifying small-molecule metabolites or lipids in complex clinical specimens such as serum and tissue samples (*10*). Two basic types of metabolomics experiments exist, either targeted or non-targeted (*11*). These experiments are typically conducted using nuclear magnetic resonance (NMR) spectroscopy and/or mass spectrometry (MS). Non-targeted metabolomics/lipidomics allows for the unbiased detection of thousands of metabolites/lipids, while targeted approaches focus on a known set of target species. For an unbiased discovery investigation of a specific disease, as in this work, non-targeted approaches are typically the first step. Non-targeted workflows lead to the generation of big data, necessitating mining methods such as machine learning. These methods are a subset of artificial intelligence that involve developing systems that can learn and improve with more experience without being explicitly programmed to do so (*12*). Combining machine learning with metabolomics and lipidomics is a powerful approach to learn about cancer biology (*13*), providing a unique opportunity for the discovery of candidate prognostic and predictive biomarkers.

Multiple studies have attempted to find metabolome or lipidome alterations associated with ovarian cancer in biofluids (*14-18*). In Gaul *et al*., using serum metabolomics, serous epithelial ovarian cancer (EOC) was discriminated from healthy controls (HC) (HC *n* = 49, EOC *n* = 46) using 16 metabolites including numerous lipids (*14*). The discrimination achieved 100% accuracy in the cohort studied using support vector machines (SVM) (*14*). Braicu and co-workers conducted a serum metabolomics study detailing profound lipid metabolism alterations (*15*). Serum samples of 147 OC patients were compared with 98 control subjects with benign ovarian tumors and non-neoplastic diseases. Improved predictive values were achieved when cancer antigen 125, the current OC clinical biomarker, was used alongside some lipid species identified in the study (*15*). Metabolomics investigations on ovarian cancer mouse models have also been conducted. Jones *et al*. performed metabolomic serum profiling for the detection of early-stage HGSC in DKO mice, identifying 18 discriminatory metabolites, including lipids in the phosphatidylethanolamine (PE), triglyceride (TG), lysophosphatidylethanolamine (LysoPE), and phosphatidylinositol (PI) classes (*19*).

Here, we present the first in-depth machine learning longitudinal analysis of the serum lipidome of a DKO HGSC mouse model using a four-pronged approach: 1) unsupervised machine learning methods and univariate statistical analyses to map global lipidome alterations, 2) hierarchical clustering analysis to identify lipidome changes in response to HGSC progression, 3) multiple machine learning algorithms with varying inductive biases to identify time-resolved HGSC evolution, and 4) Kaplan-Meier estimates and Restricted Mean Survival Times analyses to find prognostic circulating lipid marker candidates.

## Results

### Research design and computational pipeline

To study HGSC development and progression, we employed DKO mice (*Dicer1* ^flox/flox^ *Pten* _flox/flox_ *Amhr2* ^cre/+^) and DKO control mice (*Dicer1* ^flox/flox^ *Pten* ^flox/flox^ *Amhr2* ^+/+^) models using high-density blood sampling (**Figure 1a**). A total of 15 mice in both groups were used for analysis. Starting from the two-month mark, blood samples were collected biweekly until humane sacrifice of the animals, or at the end of the study at 46 weeks. This longitudinal design resulted in 221 and 238 blood samples collected for DKO and DKO control mice, respectively. As expected, DKO mice had a shorter lifespan than DKO control mice, as shown by the Kaplan-Meier (**Figure S1a**) and the Nelson-Aalen (**Figure S1b**) estimate curves. Furthermore, the restricted mean survival time difference (ΔRMST) between DKO and DKO control mice was about three weeks (**Figure S1c**).

**Figure 1.**
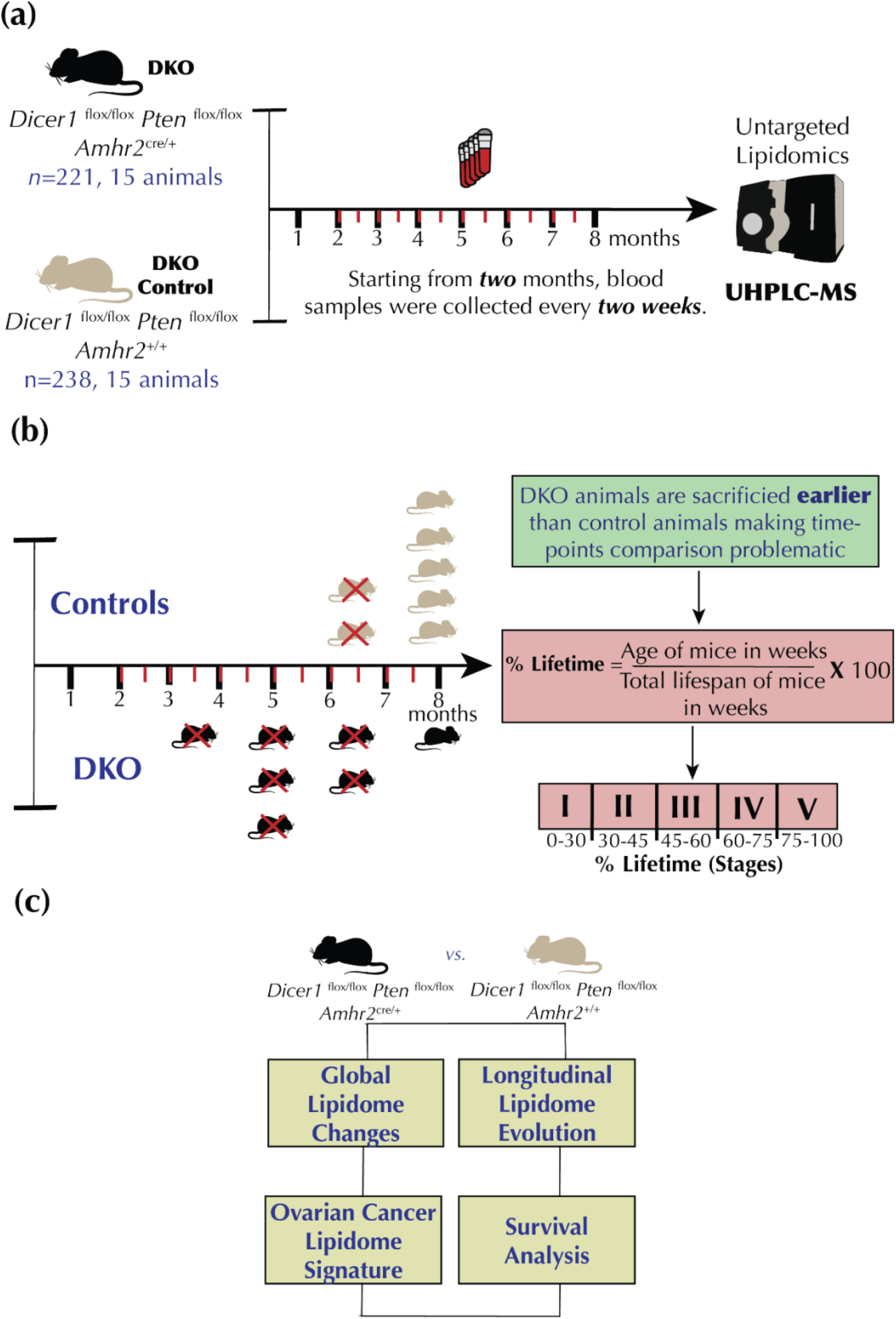
Blood sampling scheme, study design, and analysis plan. **(a)** Blood samples were collected every two weeks, starting at the two-month mark. Lipidomics experiments were conducted using ultra-high performance liquid chromatography mass spectrometry (UHPLC-MS). **(b)** Conversion of the mice age in weeks to percentage lifetime makes lipidomic comparisons effective. **(c)** Computational analysis plan.

Given the time-course data misalignment, each time point was converted to a “percentage lifetime” variable to align the dataset (**Figure 1b**). The percentage lifetime was computed by taking the percentage of the age of each mouse in weeks normalized by the total lifespan of the mouse (or age of the mice) at the last time point of blood collection (see Methods). Percent lifetimes were binned into five stages, which we named the “lifetime stage”. 0-30% lifetime was named as lifetime stage I (DKO *n* = 28, DKO control *n* = 34), 30-45% lifetime was lifetime stage II (DKO *n* = 41, DKO control *n* = 45), 45-60% lifetime was lifetime stage III (DKO *n* = 43, DKO control *n* = 42), 60-75% lifetime was lifetime stage IV (DKO *n* = 41, DKO control *n* = 45), and 75-100% lifetime was lifetime stage V (DKO *n* = 68, DKO control *n* = 72). Where *n* refers to the number of time points present in each lifetime stage.

The longitudinal lipidomic dataset was then investigated to (1) identify global lipidome alterations between DKO and DKO control mice within these lifetime stages, (2) investigate the longitudinal lipidome evolution in response to HGSC progression, (3) identify lipidome signatures for each of the five lifetime stages *via* supervised ML, and (4) identify prognostic circulating candidate biomarkers *via* survival analysis (**Figure 1c**).

### Global lipidomic changes in the DKO HGSC model

In-depth lipidomic profiling of all 459 serum samples was carried out using reverse-phase (RP) ultra-high performance liquid chromatography-mass spectrometry (UHPLC-MS). A total of 17,293 and 4,414 features (de-adducted and de-isotoped *m/z*, retention time pairs) were extracted from the RP UHPLC-MS dataset in the positive and negative ion modes, respectively. After data curation and structural annotation, 1070 lipids were identified by matching to an in-house lipid MS^2^ database. The classes of lipids detected included triacylglycerols (TG), fatty acids (FA), hexosylceramides (HexCer), lysophosphatidylcholines (LPC), lysophosphatidylethanolamines (LPE), phosphatidylcholines (PC), ether phosphatidylcholines (PC-O), phosphatidylethanolamines (PE), ether phosphatidylethanolamines (PE-O), phosphatidylinositols (PI), ceramides (Cer), sterols, and sphingomyelins (SM). **Figure 2a** shows fold changes (Log_2_FC [DKO/control]) for all 1070 annotated lipids and time points combined, indicating significant lipidome remodeling. Of the 1070 compounds annotated, 87 lipids (**Table S1**) had corrected *P*-values lower than 0.05 (Welch’s *T*-test, Benjamini-Hochberg (BH) correction *q*-value < 0.05). Some of the top-most altered lipids included HexCer(d34:1), PE(O-37:6), PE(O-36:6), and FA(14:1) (**Figure 2b**).

**Fig 2.**
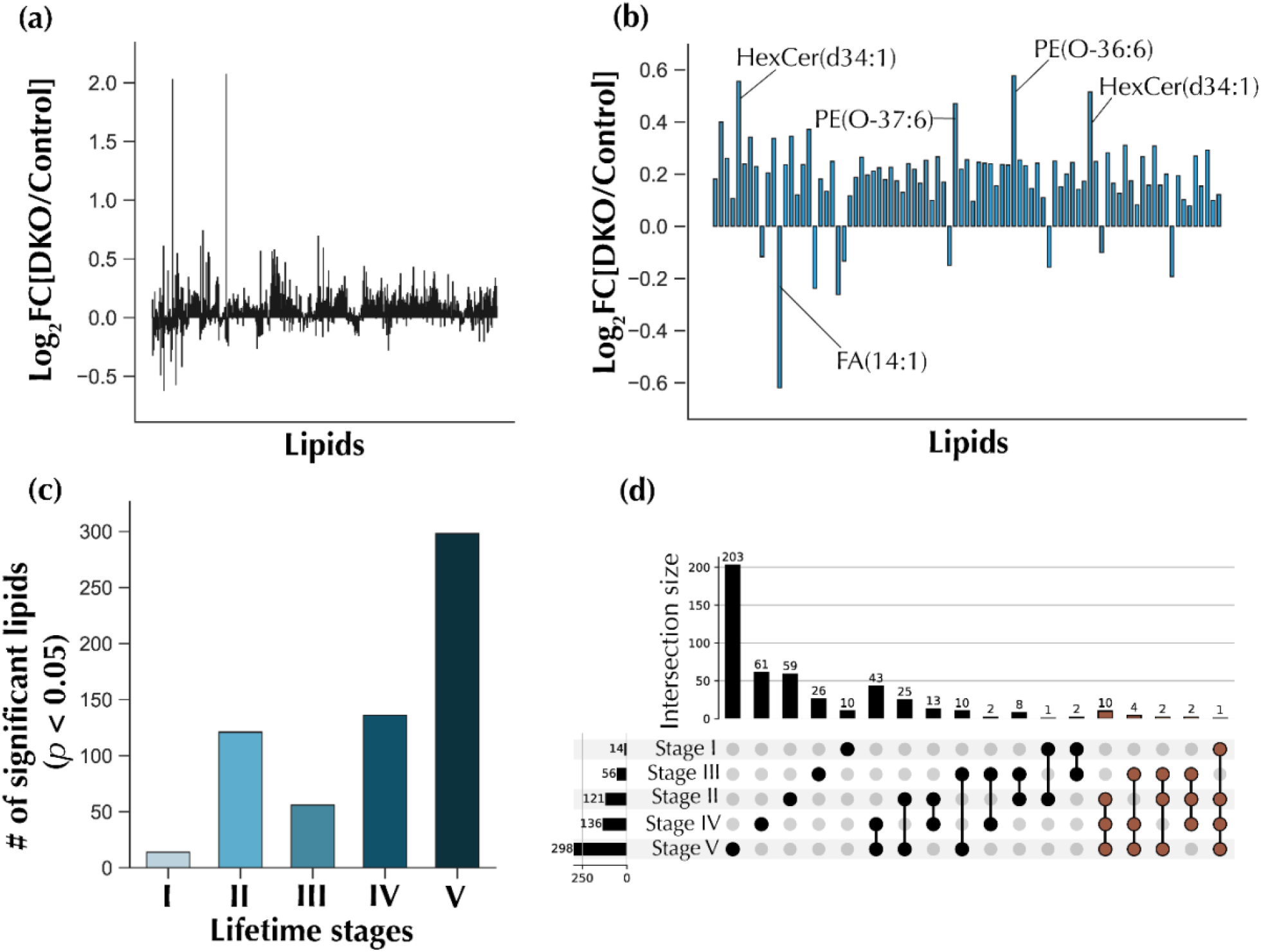
Global lipidomic changes observed upon HGSC progression. **(a)** Overall fold changes for all annotated features, for all time points combined. **(b)** Fold changes for 87 significant lipid features (Welch’s *T*-test, Benjamini-Hochberg correction *q*-value < 0.05) for all time points combined. **(c)** The number of significant lipidomic features (Welch’s T-test *p*-value < 0.05) for each lifetime stage. **(d)** Upset plot showing overlapping significant lipids in various lifetime stages. Sets containing lipid features present in at least three lifetime stages are colored brown.

To investigate global differences between DKO and DKO control mice, the 87 significant lipids were used to conduct unsupervised learning for all combined time points. PCA (**Figure S2a**), kernel PCA (**Figure S2b**), t-SNE (**Figure S2c**), and uMAP (**Figure S2d**) analyses were conducted; however, clear group clustering was unsuccessful. We also investigated time-resolved lipidome remodeling in DKO *vs*. DKO control mice through standard univariate analysis. For each lifetime stage, the number of significant lipid features was identified (Welch’s *T*-test *P*-value < 0.05). Fourteen lipids were significant in lifetime stage I, 121 in lifetime stage II, 56 in lifetime stage III, 136 in lifetime stage IV, and 298 in lifetime stage V (**Figure 2c**). There was a progressive increase in the number of significantly altered lipids as HGSC advanced, except for the observed decrease from lifetime stage II to III. This overall temporal trend seems to mimic HGSC evolution in humans where the disease evolves from an asymptomatic early stage with only minimal metabolic changes to being more easily detectable at more advanced stages where profound metabolic changes are expected. A breakdown for the significant lipids common across stages is presented in the upset plot in **Figure 2d**. A total of 71.4 % of the lipids were unique to lifetime stage I, 48.8 % to stage II, 46.4 % to stage III, 44.8 % to stage IV, and 68.1 % to stage V. Furthermore, a total of 19 serum lipids were found to be significantly altered in at least three of the five lifetime stages (**Table S2**). Of these, 68.4 % were PC or PC-O, making these the most upregulated lipid classes based on univariate time-resolved analysis.

### Lipidome alterations in response to ovarian cancer progression

Taking advantage of the granularity of our longitudinal RP UHPLC-MS dataset, we investigated lipidome changes associated with OC progression by identifying lipid trajectory clusters and calculating pairwise correlations between lipids in each cluster (**Figure 3, Table 1**). The dataset consisting of 87 significant lipids (Welch’s *T*-test, BH *q*-value < 0.05, DKO *vs*. DKO mice) was used for this analysis. To study the temporal evolution of these lipid alterations, time-resolved average lipid abundances in DKO and DKO control mice were computed. Using fold changes between the average lipid abundances (Log_2_[DKO/control])), hierarchical clustering was used to identify four main lipid trajectory clusters (A-D). In cluster A, the lipid fold changes increased in DKO mice from lifetime stage I to II, decreased from II to III, and then spiked back up in V. Similar temporal trends were observed for cluster B lipids. However, in cluster C, lipids increased from lifetime stage I to II, decreased from II to III, and increased back from III to IV, followed by a mostly slight downward trend from lifetime stage IV to V. Finally, cluster D lipids had a relatively mild temporal change from lifetime stage I to IV, with a sharp increase from IV to V (**Figure 3a-b**). A correlation network graph for these clusters is presented in **Figure 3c**, showing the connectivity of related and the same lipid classes. A common characteristic of clusters A, B, and C was an increase of the specific lipids in DKO mice from lifetime stage I to II, followed by a decrease in from stage II to III. These clusters were mostly composed of ether-linked and ester phospholipids such as PC, PC-O, PE, PE-O, and LPE. Of these lipid classes, PC and PC-O were the most represented, with 53.8% in cluster A, 100% in cluster B, and 88.8% in cluster C. On the other hand, sphingolipids classes such as HexCer and Cer comprised 79% of all cluster D lipid species. Significant serum lipidome rewiring was apparent with disease progression as shown by clustering analysis, with mostly PC and PC-O being perturbed at early stages and HexCer and Cer at advanced stages.

**Table 1.** Annotations for lipid clusters associated with ovarian cancer progression. Proposed lipid annotation, experimental *m/z* value, chromatographic retention time (RT) in minutes (min), and main adduct type detected are shown. DG: Diacylglycerols, TG: Triacylglycerols, FA: Fatty acids, HexCer: Hexosylceramides, LPC: Lysophosphatidylcholines, LPE: Lysophosphatidylethanolamines, PC: Phosphatidylcholines, PC-O: Ether phosphatidylcholines, PE: Phosphatidylethanolamines, PE-O: Ether phosphatidylethanolamines, PI: Phosphatidylinositols, Cer: Ceramides, and SM: Sphingomyelins.

| ID | Lipids | Adduct | RT [min] | Mass Error (ppm) | Experimental $m/z$ |
| --- | --- | --- | --- | --- | --- |
| Cluster A |  |  |  |  |  |
| 4260 | HexCer(d42:2) | $[M+CH_3COOH-H]^-$ | 6.50 | 3.93 | 868.6917 |
| 2626 | PC(20:0_20:4) | $[M+H_2CO_2-H]^-$ | 5.20 | 3.13 | 882.6257 |
| 1876 | PC(O-16:0_16:0) | $[M+CH_3COOH-H]^-$ | 5.17 | 4.31 | 778.6001 |
| 2412 | PC(O-18:1/20:3) | $[M+CH_3COOH-H]^-$ | 5.03 | 3.36 | 854.6309 |
| 2587 | PC(O-18:1_22:6)<br>and PC(O-22:7_18:0) | $[M+CH_3COOH-H]^-$ | 4.33 | 3.31 | 876.6153 |
| 1650 | PE(16:0_20:4) | $[M-H]^-$ | 4.35 | 4.46 | 738.5122 |
| 1651 | PE(16:0_20:4) | $[M-H]^-$ | 4.17 | 4.62 | 738.5113 |
| 1623 | PE(O-15:1_22:5) | $[M-H]^-$ | 4.35 | 4.71 | 734.5164 |
| 10366 | TG(60:12) | $[M+NH_4]^+$ | 7.56 | 2.82 | 968.7729 |
| 6813 | HexCer(d18:1_24:1) | $[M+H]^+$ | 6.54 | 4.42 | 810.6853 |
| 6839 | PC(38:3) | $[M+H]^+$ | 4.95 | -1.03 | 812.6155 |
| 7604 | PC(41:6) | $[M+H]^+$ | 5.04 | 1.81 | 848.6179 |
| 7565 | PC(41:7) | $[M+H]^+$ | 5.26 | -2.04 | 846.5990 |
| Cluster B |  |  |  |  |  |
| 1870 | PC(16:0_16:0) | $[M+H_2CO_2-H]^-$ | 4.26 | 3.38 | 778.5629 |
| 2050 | PC(16:0_18:0) | $[M+H_2CO_2-H]^-$ | 5.45 | 4.03 | 806.5949 |
| 1679 | PC(16:0_18:2) | $[M+H_2CO_2-H]^-$ | 4.14 | -0.76 | 802.5597 |
| 2411 | PC(O-18:1_20:3) | $[M+CH_3COOH-H]^-$ | 4.52 | -1.85 | 854.6264 |
| 2400 | PC(O-18:1_20:4)<br>and PC(O-16:0_22:5) | $[M+CH_3COOH-H]^-$ | 4.52 | 3.18 | 852.6151 |
| Cluster C |  |  |  |  |  |
| 886 | LPE(20:0) | $[M-H]^-$ | 2.27 | 4.47 | 508.3431 |
| 6573 | PC(37:3) | $[M+H]^+$ | 4.80 | 2.48 | 798.6027 |
| 7105 | PC(39:4) | $[M+H]^+$ | 5.26 | 0.83 | 824.6170 |
| 5618 | PC(O-32:0) | $[M+H]^+$ | 5.22 | 1.41 | 720.5911 |
| 5604 | PC(O-32:1) | $[M+H]^+$ | 4.48 | 1.61 | 718.5756 |
| 6493 | PC(O-38:5) | $[M+H]^+$ | 4.56 | -0.28 | 794.6060 |
| 7022 | PC(O-40:6) | $[M+H]^+$ | 5.26 | 0.73 | 820.6220 |
| 7023 | PC(O-40:6) | $[M+H]^+$ | 4.62 | 2.81 | 820.6237 |
| 6984 | PC(O-40:7) | $[M+H]^+$ | 4.40 | 0.10 | 818.6059 |
| **Cluster D** |  |  |  |  |  |
| 1111 | Cer(d33:1) | $[M+CH_3COOH-H]^-$ | 4.69 | 4.80 | 582.5131 |
| 966 | Cer(d34:1) | $[M-H]^-$ | 4.70 | 2.96, | 536.5064, |
| 1149 | Cer(d34:1) | $[M+CH_3COOH-H]^-$ | 4.72 | 4.28 | 596.5285 |
| 1217 | Cer(d40:1) | $[M-H]^-$,<br>$[M+CH_3COOH-H]^-$ | 6.82 | 4.62, 4.38 | 620.6014,<br>680.6229 |
| 1297 | Cer(d42:2) | $[M-H]^-$ | 6.79 | 4.73, | 646.6174, |
| 1290 | Cer(d42:3) | $[M-H]^-$ | 6.45 | 4.27 | 644.6011 |
| 1504 | Cer(d42:3) | $[M+CH_3COOH-H]^-$ | 6.46 | 4.19 | 704.6228 |
| 1473 | HexCer(d34:1) | $[M-H]^-$ | 4.32 | 3.17 | 698.5611 |
| 1761 | HexCer(d34:1) | $[M+CH_3COOH-H]^-$ | 4.09 | 4.21 | 758.5819 |
| 1762 | HexCer(d34:1) | $[M+CH_3COOH-H]^-$ | 4.32 | 2.37 | 758.5805 |
| 2078 | HexCer(d42:1) | $[M-H]^-$ | 6.83 | 4.50 | 810.6857 |
| 2532 | HexCer(d42:1) | $[M+CH_3COOH-H]^-$ | 6.83 | 2.53 | 870.7061 |
| 2065 | HexCer(d42:2) | $[M-H]^-$ | 6.40 | 3.76 | 808.6698 |
| 2522 | HexCer(d42:2) | $[M+CH_3COOH-H]^-$ | 6.41 | 3.83 | 868.6916 |
| 2415 | HexCer(d42:2) | $[M+H_2CO_2-H]^-$ | 6.40 | 3.70 | 854.6758 |
| 2557 | SM(d42:1) | $[M+CH_3COOH-H]^-$ | 7.16 | 3.95 | 873.7100 |
| 7815 | PC(42:8) | $[M+H]^+$ | 4.08 | 1.32 | 858.6018 |
| 9401 | TG(56:9) | $[M+NH_4]^+$ | 7.59 | 0.26 | 918.7547 |
| 10226 | TG(58:9) | $[M+NH_4]^+$ | 7.65 | 5.36 | 946.7908 |

**Fig 3.**
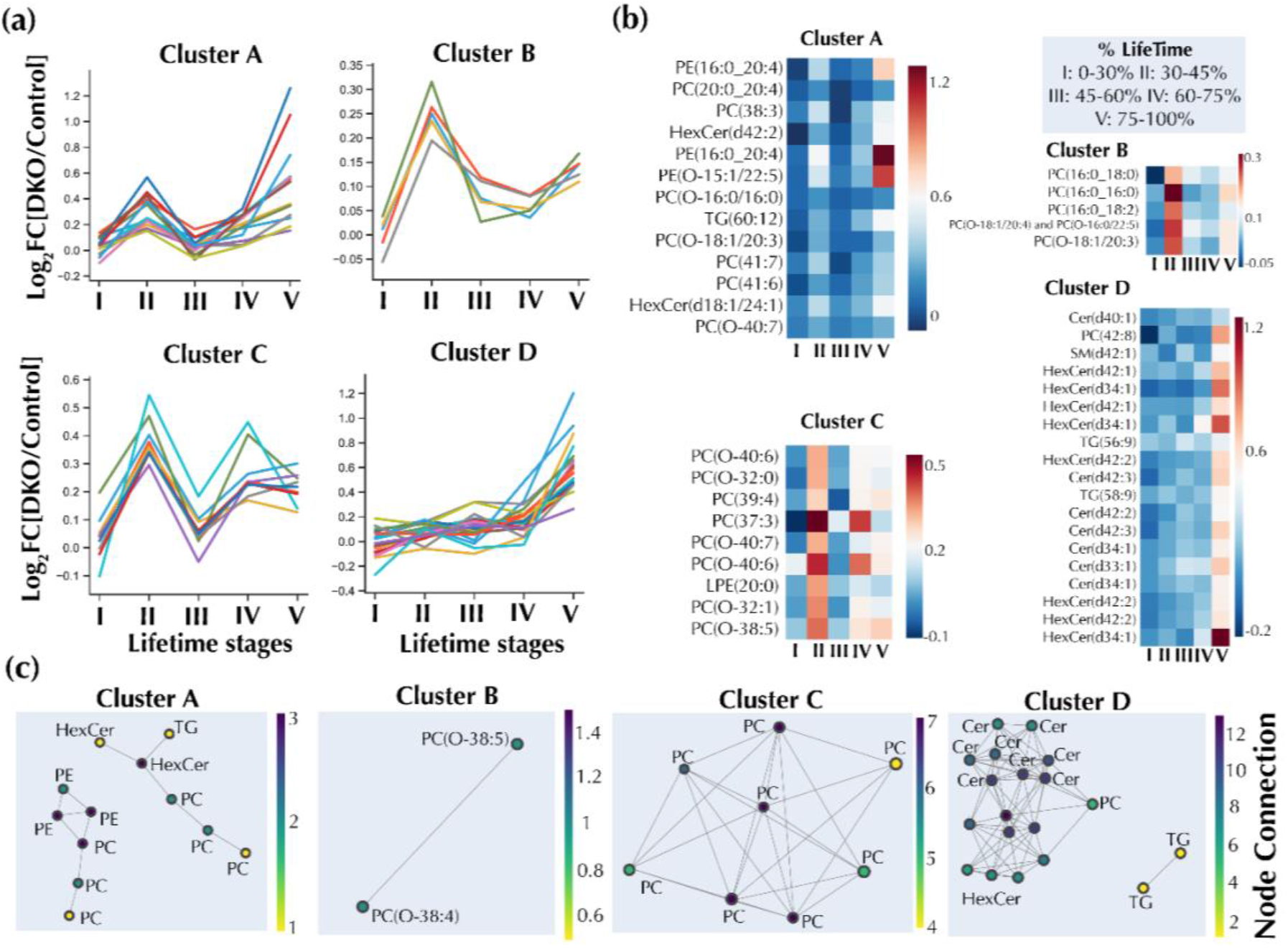
Lipidome changes in response to ovarian cancer progression. **(a)** Hierarchical clustering analysis shows the grouping of lipidome trajectories into four types of clusters. **(b)** Longitudinal lipid changes for the selected clusters indicating fold changes. **(c)** Network graph for the clusters shown in (a). Nodes represent lipids, while the links connect nodes with a high Pearson’s correlation (r ≥0.5).

### Time-resolved machine learning discriminates tumor stages of HGSC in DKO mice

We subsequently employed in-depth machine learning (ML) to further characterize the five-lifetime stages. The feature selection strategy in the ML computational pipeline (**Figure 4a**) led to the selection of five lipid features for lifetime stage I, 25 for lifetime stage II, 18 for lifetime stage III, 24 for lifetime stage IV, and 42 for lifetime stage V (**Table S3**). After feature selection, five ML algorithms, including logistic regression, random forests (RF), *k*-nearest neighbors (*k*-NN), support vector machine (SVM), and a voting classifier composed of the four prior ML algorithms were used to discriminate DKO from DKO control mice within each of the lifetime stages (**Figure 4a**). ML algorithms were trained under fivefold cross-validation conditions, while a separate test set was used for testing purposes. Detailed ML prediction results are presented in **Table S4**. For lifetime stage I (training set *n* = 43, test set *n* = 19), RF, *k*-NN, and a voting classifier gave the best receiver operating curve area under the curve (ROC AUC) test set score of 0.80 (**Figure 4b** and **g**). For lifetime stage II (training set *n* = 60, test set *n* = 26), RF gave the highest ROC AUC test set score of 0.70 (**Figure 4c** and **g**). For lifetime stage III (training set *n* = 59, test set *n* = 26), logistic regression had the highest ROC AUC test set score of 0.85 (**Figure 4d** and **g**). For lifetime stage IV (training set *n* = 60, test set *n* = 26), RF gave the highest ROC AUC test set score of 0.66 (**Figure 4e** and **g**), and finally, for lifetime stage V (training set *n* = 98, test set *n* = 42), SVM gave the highest score of 0.75 (**Figure 4f** and **g**).

**Fig 4.**
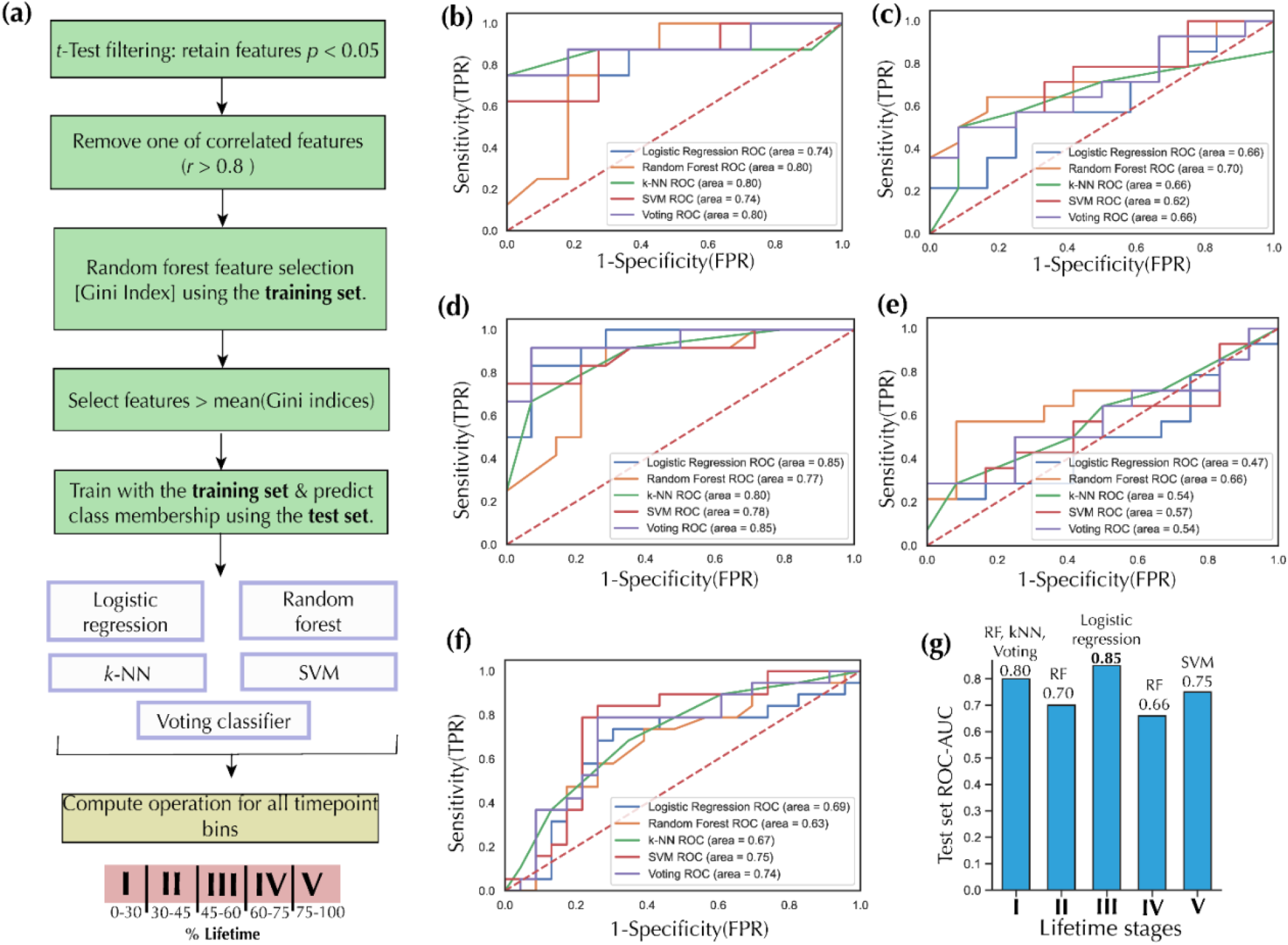
Discriminating DKO from DKO control mice *via* machine learning. **(a)** Machine learning pipeline. The pipeline starts with a *t*-test filtering method for each of the five ML tasks: lipid features with less than 0.05 *p*-value (Welch’s *t*-test *p* < 0.05) were selected. Next, one of two lipid features with a high Pearson’s correlation score (*r* > 0.8) was removed from the dataset to avoid unnecessary redundancies. Finally, lipid features with a Gini index greater or equal to the mean of all Gini indices were selected for training and testing purposes. ROC-AUC test set for DKO classification for **(b)**, lifetime stage I **(c)**, lifetime stage II **(d)**, lifetime stage III **(e)**, lifetime stage IV **(f)**, and lifetime stage V. **(g)** The best ROC-AUC scores for each lifetime stage. TPR: True positive rate, FPR: False positive rate, k-NN: k-Nearest Neighbors, RF: Random Forests, SVM: Support Vector Machine, Voting: Voting Ensemble Classifier.

Given that early detection of ovarian cancer is important for improving clinical outcomes, an AUC value of 0.80 for the first-lifetime stage (0-30%) suggests the possibility of early detection of OC *via* serum lipidomics, should the lipids in the panel also show significant alterations in humans. The discriminant lipids included a medium-chain fatty acid, 3-hydroxyphenyl-valerate, and four phospholipids: PE(O-34:3), PC(17:0_18:2), PC(38:6), and PE(O-16:1_20:5) (**Figure 5a** and **Table S3**). Furthermore, the highest AUC value for the five lifetimes was 0.85 for lifetime stage III (45-60%); the selected discriminant lipid features included ester phospholipids PC(18:0_18:0), PC(16:0_20:4), PC(18:0_20:4), PC(18:0_22:4), PC(37:6), and PI(18:1_20:4), ether phospholipids PE(O-18:0_18:2) and PC(O-38:6), ceramides Cer(d33:1), Cer(d41:2), Cer(d45:1), cerebrosides HexCer(d38:0-OH) and HexCer(d40:0) or HexCer(t42:0-OH), a fatty acid FA(18:2), a glycerol ester, TG(18:0_18:1_18:2), prostaglandin A1, and a pyrimidine derivative (**Figure 5c, Table S3**). Other selected lipid markers for lifetime stages II, IV, and V are shown in **Figures 5b, d, e**, and **Table S3**. A summary of the lipid categories represented in each of the ML discriminant panels is given in **Figure 5f**. Phospholipids were the most represented category in all the five lipid discriminant panels. Of all the phospholipid classes, PC and PC-O were the most abundant species. The least represented lipid category was steroid lipids, with just one cholesterol derivative selected in the lifetime stage V (75-100%) panel. Furthermore, of all the lipids selected as markers, only phospholipids and fatty acyls (composed mostly of fatty acids) were selected in all the lifetime stages. In summary, the early progression of OC was marked by increased levels of phospholipids, notably PC and PC-O while, in contrast, later stages were marked by more diverse lipids alterations, including sphingolipids, fatty acyls, glycerolipids, steroid lipids, and phospholipids. Apart from phospholipids, sphingolipids were the most represented lipid category at stages IV and V, consisting of mostly HexCer, Cer, and SM (**Figure 5f**). These results agree with the lipid trajectory clustering results discussed earlier.

**Fig 5.**
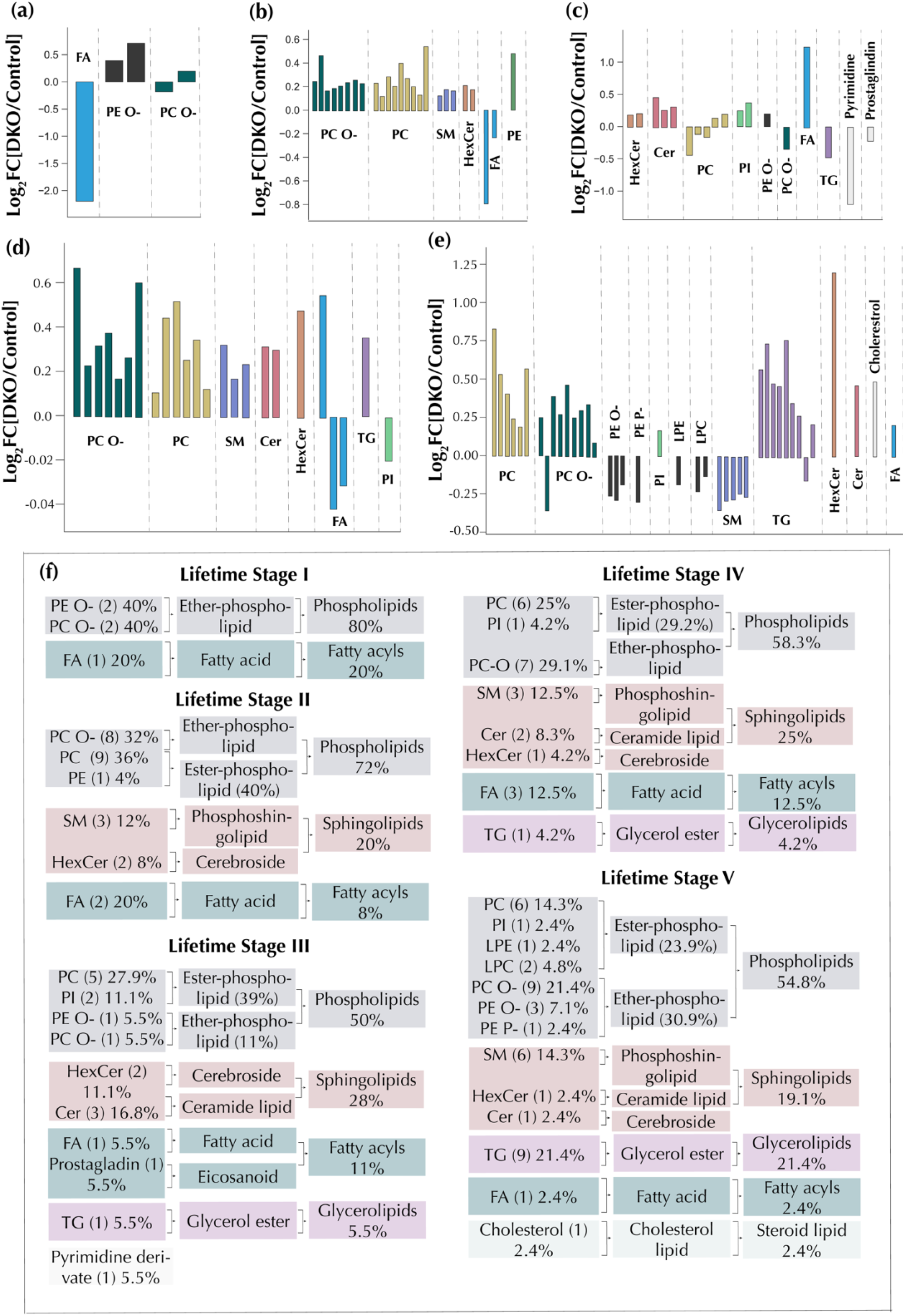
Discriminant lipids for each of the five lifetime stages. **(a)** Lifetime stage I: 0-30% lifetime. **(b)** Lifetime stage II: 30-45% lifetime. **(c)** lifetime stage III: 45-60% lifetime **(d)** Lifetime stage IV: 60-75% lifetime. **(e)** Lifetime stage V: 75-100% lifetime. **(f)** frequency of lipid classes, groups, and categories in the discriminant lipid panels. TG: Triacylglycerols, FA: Fatty acids, HexCer: Hexosylceramides, LPC: Lysophosphatidylcholines, LPE: Lysophosphatidylethanolamines, PC: Phosphatidylcholines, PC-O: Ether phosphatidylcholines, PE: Phosphatidylethanolamines, PE-O: Ether phosphatidylethanolamines, PI: Phosphatidylinositols, Cer: Ceramides, and SM: Sphingomyelins

### Prognostic circulating lipids in DKO mice

Because prognostic makers are useful in providing information on the likely health outcome of cancer patients, we employed survival analysis methods to investigate lipid species predictive of the course of OC in DKO mice. First, candidate lipids were selected by comparing all 1070 lipid features in DKO lifetime stages II – V with DKO lifetime stage I. Lipids features with *p*-values < 0.05 (Welch’s *T*-test) and at least one fold change (log_2_FC, DKO lifetime stage II-V *vs*. DKO stage I) were selected, resulting in a set of ten different lipids in DKO lifetime stages I *vs*. II (**Figure 6a**), 56 in I *vs*. III (**Figure 6b**), 68 in I *vs*. IV (**Figure 6c**), and 29 in I *vs*. V (**Figure 6d**). A breakdown of overlapping and unique lipid features in these subsets is given in the upset plot in **Figure 6e**. A total of 12 lipids were present in at least three sets from various lifetime pair comparisons. These lipids were selected as prognostic candidates (**Figure 6e**). Furthermore, the 19 lipid features found to be differential in at least three of the five lifetime stages (**Figure 2d**) were also selected as candidate prognostic lipids. All fifteen DKO animals were binned into two groups based on a median split using all 31 candidate prognostic lipids. A DKO ‘Low’ group was built from mice with lipid abundances lower than or equal to the median of the relative abundances of the selected lipids, while mice with abundances greater than the median were bundled into a DKO ‘High’ group. Three lipid species of the 31 lipid candidates had a statistically significant difference in their Kaplan Meier (KM) curves *via* the log-rank test. These included PC(39:4) (*p*-value = 0.003, **Figure 6f**), PC(37:2) (*p*-value = 0.02, **Figure 6g**), and PC(40:7) (*p*-value = 0.008, **Figure 6h**). Of the 3 prognostic lipids, PC(39:4) had the strongest prognostic effects with an ΔRMST of 10.96, followed by PC(40:7) (ΔRMST = 9.35), and then PC(37:2) (ΔRMST = 7.75) (**Figure S3**). All the prognostic circulating lipids had elevated levels in DKO mice compared to DKO control mice for all time points combined (**Figure 6h**).

**Fig 6.**
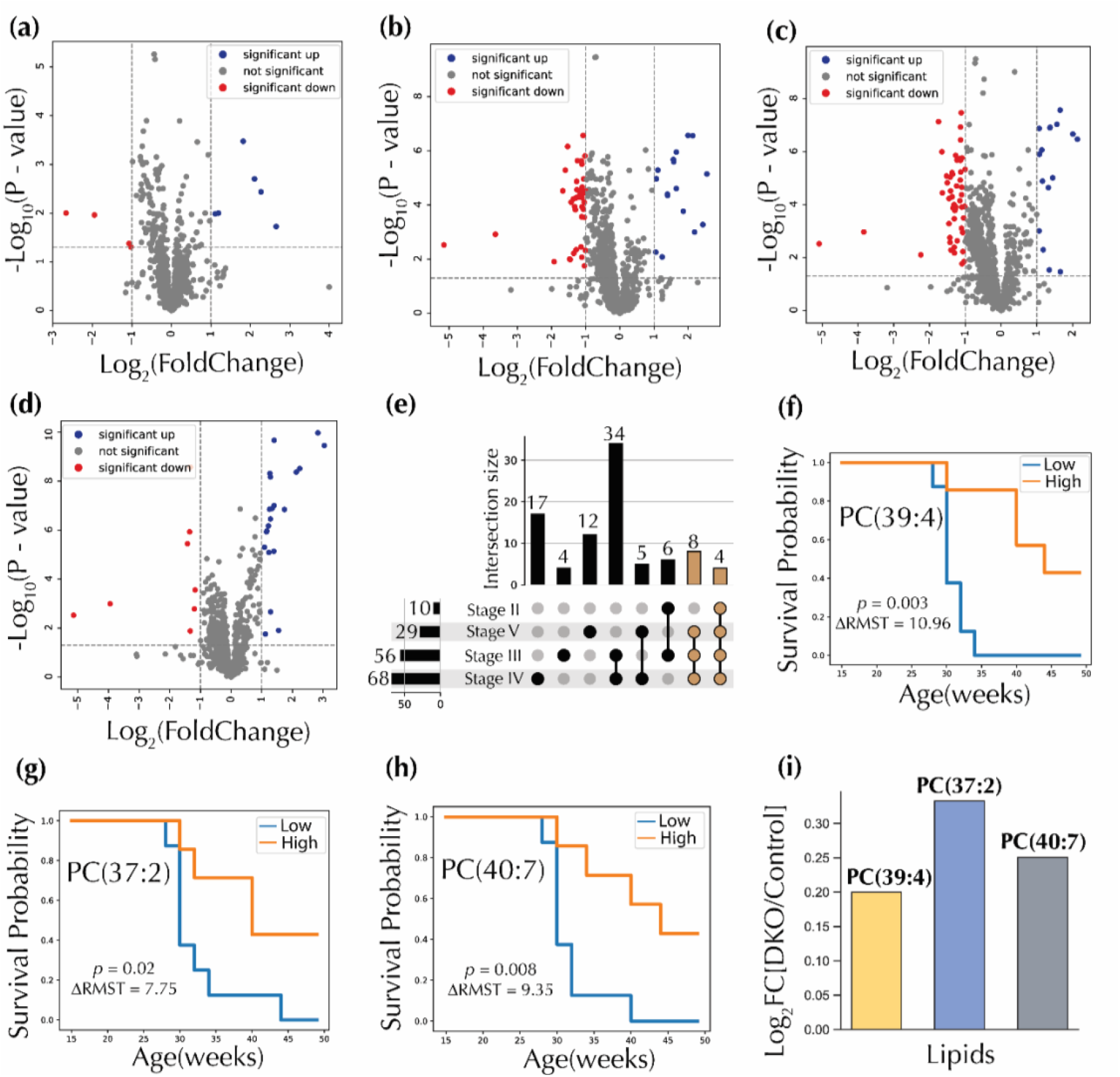
Prognostic circulating lipid candidates. Volcano plots comparing DKO lifetime stage I with **(a)** DKO lifetime stage II, **(b)** DKO lifetime stage III, **(c)** DKO lifetime stage IV and **(d)** DKO lifetime stage V. *P*-values for volcano plot analysis were calculated using Welch’s *T*-test. **(e)** Upset plot showing the intersection of the various groups of significant lipids selected from volcano plots. Lipids present in at least three sets were colored brown. Kaplan-Meier survival curves for **(f)** PC(39:4), PC(37:2) and **(h)** PC(40:7). *P*-values were computed with the Log rank test. **(i)** Selected prognostic circulating lipids. PC: Phosphatidylcholines FC: Fold changes. ΔRMST: differences in restricted mean survival times.

## Discussion

Given that most metabolomic cancer studies are based on a snapshot of the metabolic process (*14-18*), it is not surprising that an understanding of the metabolic pathogenesis of HGSC remains elusive. In this study, we performed nontargeted serum lipidomics of DKO mice, an ovarian HGSC mouse model. We examined the temporal interplay of serum lipids in ovarian HGSC progression. Ovarian HGSC originates in the fallopian tube where fallopian tube epithelial (FTE) cells may be transformed into serous tubal intraepithelial carcinoma (STIC) lesions. STIC metastasize into the ovary and then to the omentum (*20*). The omentum, an extensive network of adipose tissue, provides a secondary metastasis hub (*21, 22*), further underscoring the importance of investigating ovarian HGSC pathogenesis through lipidome alterations. Reassuringly, our study identified similarly altered lipids as a previous study at a fixed time point (*19*), validating the experimental approach applied here. As expected, and given the pathogenesis of HGSC (*20*), significant lipid alterations were evident from the data analysis performed when all time points were combined. The most altered lipid classes at a global level included sphingolipids and phospholipids, with the general trend showing that the number of significant lipids for each lifetime stage increased as ovarian HGSC progressed. PC and PC-O were the most perturbed lipid classes, following perturbations shown in previous metabolomic studies (*23*).

### Phospholipids

Phospholipids, specifically ether and ester phospholipids, are by far the predominant lipid classes present in clusters A-C of the temporal trend analyses conducted in this study, with PC and PC-O being the key lipid families. This finding is not surprising, as PC comprise approximately 40-50% of all total cellular phospholipids (*24*). Furthermore, cancer cells require increased generation and maintenance of cellular membranes, largely composed of phospholipids (*25*). Iorio *et al*. reported the activation of phosphatidylcholine-cycle enzymes in human epithelial ovarian cancer (EOC) cells (*26*). In that study, the authors reported increased phosphocholine (Pcho) levels and upregulation of choline kinase (ChoK)-mediated phosphorylation, providing a plausible explanation for the observed increase in PC levels, particularly for the progression from lifetime stage I to II in clusters A-C. These data strongly suggest upregulation of the Kennedy pathway (*27*), with a predominance of PC generation. Altered PC levels in ovarian cancer have been previously reported in human studies (*28*) and in an ovarian cancer mouse model (*23*). This temporal trend for phospholipids agrees with the discriminant lipids selected for DKO classification tasks for all lifetime stages (Figure 5f). PCs and PC-Os comprise most of the lipids selected for classification within lifetime stage II. In addition, phospholipids have the highest percentage of discriminant lipids at all lifetime stages, with a decreasing proportion as HGSC progresses. This finding suggests that phospholipids may play lesser roles in advanced HGSC. In addition, three PC species (PC(39:4), PC(37:2), and PC(40:7)) were identified as potential prognostic circulating lipids.

Of all discriminant lipids identified, most phospholipids species increased, while a few decreased, such as LPE and LPC. LPC perturbations have been reported in an ovarian cancer human study (*28*) and LPE species have been suggested as early-stage ovarian cancer biomarkers in another human study (*14*). In a study of the triple knock out (TKO) HGSC mouse model, LPE and LPC were likewise altered (*23*). In our study, LPE(18:1), LPC(20:4/0:0), and LPC(20:5/0:0) were selected as discriminant lipids for lifetime stage V, with decreased levels in DKO mice. LPC and LPE are the first step in Land’s cycle, the biochemical pathway involved in the remodeling of PC and PE (*29*). LPC and LPE are mainly derived from partial hydrolysis of PC and PE, respectively, *via* phospholipase A_1_ and A_2_ (PLA_1_ & PLA_2_) (*30*). Decreased relative abundances of these lipid classes at lifetime stage V can be explained by the sustained upregulation of PC and PE. Indeed, longitudinal lipidome analysis of the TKO mouse model showed that most LPC species were lower in abundance and most PC species much higher in HGSC (*23*). Furthermore, in a large-scale profiling study of metabolic dysregulation in human ovarian cancer, LPC and LPE were reported to be elevated in localized epithelial ovarian cancer (EOC) and downregulated in metastatic EOC (*31*). These results align with findings for lifetime stage V for LPE and LPC.

Another class of phospholipids that emerged as important were the phosphatidylinositols (PI). These lipids are the central actors in the PI and PIP_2_ cycles underpinning several mammalian cell signaling pathways (*32*). There, PI is converted into phosphatidylinositol-4-phosphate (PI4P), which is further converted into phosphatidylinositol-4,5-bisphosphate (PIP_2_) *via* various phosphokinases. PIP2, on the other hand, is a component of the phosphatidylinositol 3-kinase (PI3K) pathway that has been extensively implicated in cancer (*33*). PI3Ks are lipid kinases that phosphorylate PIP2 at the 3-OH inositol group to yield phosphatidylinositol 3,4,5-trisphosphate (PIP3). PIP3 activates the serine/threonine protein kinase, which plays a key role in carcinogenesis (*33*). The perturbation of PI levels in HGSC can be rationalized by increased phosphatidylinositol 3-kinase (PI3-kinase) activity, due to the increased copy numbers of the p110α catalytic subunit of the enzyme in ovarian cancer (*34*). This altered signaling pathway has been linked to cell proliferation (*35*), glucose metabolism (*36*), and various types of oncogenic transformations (*37*). In addition, alteration of PI levels has been reported in a DKO lipidomic study (*19*), and proposed as a potential trait of early-stage OC in humans (*14*).

### Sphingolipids

Cluster D in the hierarchical clustering temporal analysis results (**Figure 3**) consists mainly of ceramides (Cer) and hexosylceramides (HexCer) with a characteristic abundance spike from lifetime stage IV to V (*i*.*e*., towards the end of the animal’s life cycle). Ceramides are essential intermediates in sphingolipid metabolism, acting as substrates for more complex sphingolipids or degradation products. For example, HexCer and sphingomyelins (SM) are derived from Cer, while SM and HexCer can be degraded to Cer by sphingomyelinases (SMAse) and cerebrosidases, respectively. Altered sphingolipid metabolism has been implicated in leukemia (*38*), hepatocellular (*39*), colorectal (*40*) and ovarian cancers (*41*). Long-chain ceramides have been identified as possible diagnostic biomarkers of human epithelial ovarian cancer (*41*). Sphingolipid metabolism has also been implicated in regulating autophagy (*42*). Autophagy’s primary role is to regulate cellular homeostasis by removing damaged organelles and aggregated proteins; however, under high-stress conditions, such as nutrition starvation, autophagy contributes to maintaining cellular functions by supplying energy to the cell (*43*). As such, in the early cancer stages, autophagy possesses an anti-carcinogenic function by attempting to maintain normal cellular operations (*43*). On the other hand, at the late stages of cancer development, autophagy confers tumor cell survival functions to counteract metabolic stress (*44*), directly explaining the temporal trends of lipids in cluster D. As such, the role of autophagy in cancer can be said to be paradoxical. Furthermore, ceramide glycosyltransferases, an enzyme class that catalyzes the formation of hexosylceramides, has been implicated in playing a role in tumor progression (*45*). Overexpression of uridine diphosphate-glucose ceramide glucosyltransferase (UGCG), the gene involved in the synthesis of glucosylceramide, has also been reported in ovarian cancer cells (*45*). The highest abundance increase for a discriminant lipid was for HexCer(d34:1) in lifetime stage V. Finally, five SM species were selected in the lifetime stage V classification task, all having low relative abundances in DKO mice *vs*. DKO controls. In contrast, cluster D lipids showed overwhelmingly increased levels of Cer and HexCer at the late stages. This metabolic trend suggests a conversion of SM to Cer via SMAse to sustain the continued proliferative effects of Cer in tumor cells.

### Fatty Acids, Triglycerides and Other Derivatives

Cancer cells can shunt energy from glucose into fatty acid synthesis (*46*), and the metabolic rearrangements are pivotal in cell signaling and tumor growth (*47*). The observed alterations in fatty acids abundances at every single lifetime stage examined are a result of this metabolic shift. Enzymes associated with lipid syntheses, such as acetyl-CoA carboxylase (ACC) and ATP-citrate lyase (ACL), are overexpressed and involved in tumorigenesis in various tumors cell types (*48-50*). Fatty acid synthase (FAS), a multi-enzyme protein whose main role is to synthesize palmitate from acetyl-CoA and malonyl-CoA, has also been found to be upregulated in ovarian cancer tissues and associated with poor disease prognosis (*51*). Furthermore, stearoyl-CoA desaturase-1 (SCD1), the enzyme that catalyzes the production of saturated fatty acids from mono-unsaturated fatty acids, is upregulated in ovarian cancer stem cells (*52*). Exogenous fatty acid metabolism also plays a role in ovarian cancer development (*46*). For instance, fatty acid binding protein (FABP4) has been identified at the interface of adipocytes and ovarian tumor cells in omental metastases (*53*). Furthermore, CD36, a member of the fatty acid transport proteins (FATP), a transmembrane transport protein that allows long-chain fatty acids into the cells, has also been implicated in breast cancer progression and metastasis (*54*). Our ML algorithm selected FA species as discriminant across all lifetime stages. Five of these were decreased in DKO mice relative to controls. These species included 3-hydroxyphenyl-valerate, FA(26:1), and FA(18:3). Changes in FA levels during tumor development most likely indicate the interplay between FA synthesis and FA cell uptake, concomitant with FA metabolism associated with the synthesis of complex lipids.

Estrogens, whose significant roles in the development and metastasis of ovarian cancer are well-documented (*55*), have been linked to increased levels of TG in mice (*56*) and humans (*57, 58*). This provides a biological link between estrogens and TG in ovarian cancer pathogenesis. Furthermore, in a metabolic study involving over a hundred thousand subjects and a ten-year follow-up period, serum TG were shown to positively correlate with gynecological (ovarian, endometrial, cervical) cancer risk (*59*). In our study, TG(60:12) was selected as one of the cluster A lipids, with levels spiking up from lifetime stage I to II, decreasing from II to III, and then increasing in stages IV and V. In addition, two triglycerides, TG(56:9) and TG(58:9), belong to cluster D lipids which have a characteristic spike from lifetime stages IV to V. For ML classification tasks, most TG played a discriminatory role in lifetime stage V, with 8 out of 9 having higher relative abundance in DKO mice. A serum metabolomics study comparing DKO mice with controls also found a triglyceride (TG 55:7) that increased in DKO mice (*19*). Triglycerides are used for energy storage, which is very much needed to support cell growth as cancer progresses. This suggests the upregulation of the monoacylglycerol and glycerol phosphate pathways.

Other selected discriminant lipids included prostaglandin A1 (PGA1), an eicosanoid. This lipid was lower in DKO mice in the third lifetime stage. Higher abundances of prostaglandin and prostaglandin D2 have been found to inhibit human ovarian cancer cell growth both *in vitro* and in mice (*60*). Similarly, A-class prostaglandins are known to have antiproliferative effects by blocking the cell cycle and activating apoptotic cascades (*61*). A cholesterol derivative was also selected as a discriminant lipid in lifetime stage V, with an increased abundance in DKO mice. Cholesterol metabolites have been linked to the promotion of tumorigenesis (*62*). Furthermore, high serum cholesterols level has been linked to increased ovarian cancer risk in a prospective study (*63*).

### Conclusions

We here present a deep temporal lipidomic study of an HGSC ovarian cancer mouse model. The main findings are summarized in **Figure 7**, pointing at numerous alterations in a variety of lipid pathways. Phospholipids were the most perturbed lipid class. They also represented the highest number of altered species at the early stages of HGSC development, pointing to cell integrity fortification processes associated with cancer progression. We also found that ceramide and hexosylceramide levels predominantly increased in DKO mice at the later stages of OC progression. It is well known that sphingolipid metabolism is linked to cancer development and progression via autophagy. In the early stages, an attempt is made to inhibit tumorigenesis; however, at later stages, those lipids assist in cancer proliferation. Furthermore, we identified sets of lipids that discriminate between DKO and DKO control mice, even at the earliest stages of disease progression. In addition, three phospholipid species were identified as circulating prognostic markers in DKO mice. These findings underscore the potential for the existence of early-stage diagnostic or prognostic lipid biomarker panels for human ovarian cancer.

**Fig 7.**
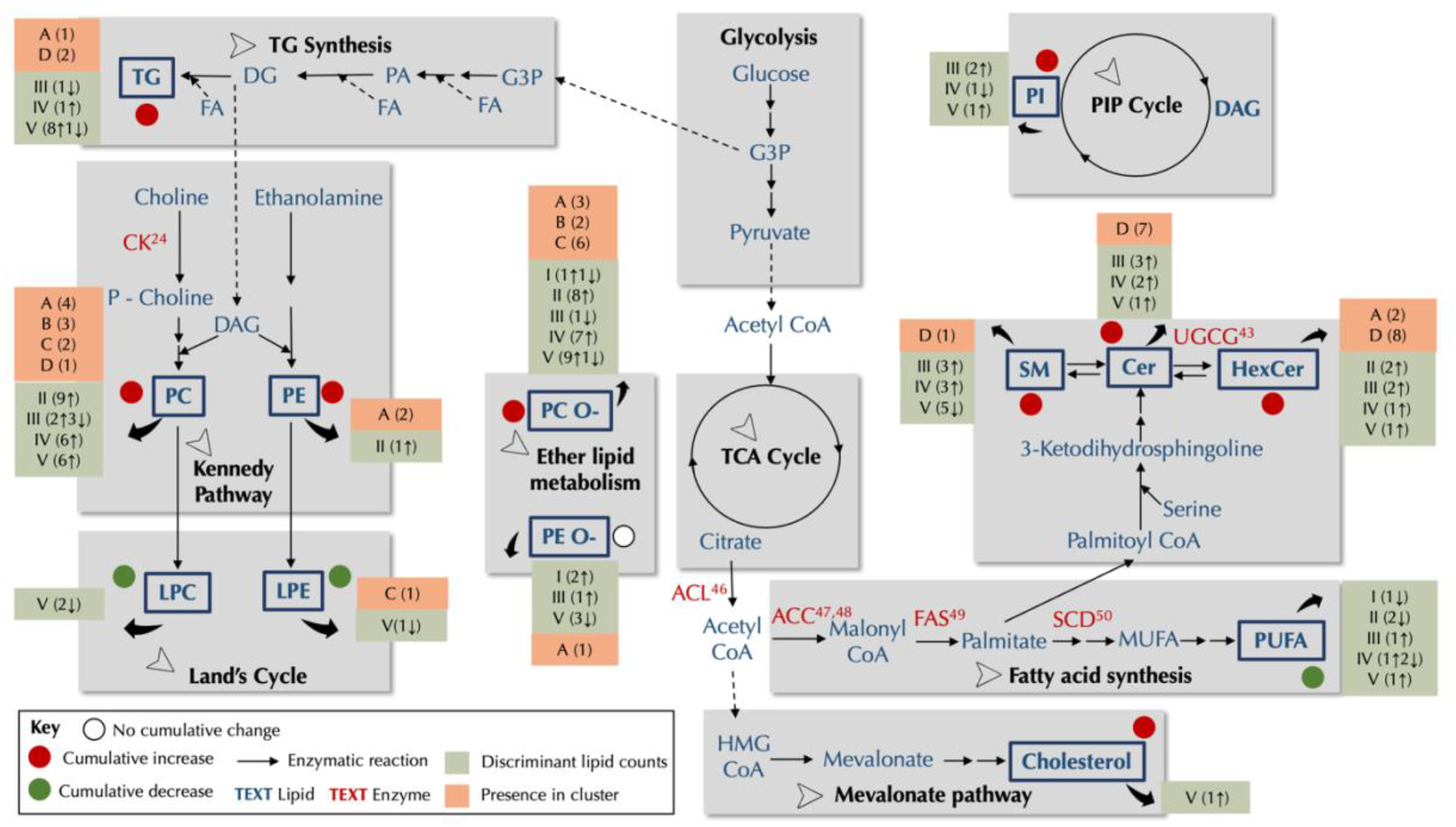
Schematic of metabolic pathways showing key metabolic alterations in the DKO mice lipidome. Lipid classes detected in the study are indicated as bolded blue text, while unbolded blue text signifies other metabolites in the metabolic pathway. Red text indicates enzymes known to be overly expressed in ovarian cancer cells or other related cancer, with the relevant references. For each detected lipid class presented, information about the cluster they belong to in the temporal trend analyses is provided, in addition to the breakdown information on discriminant lipids selected by ML algorithms. A red circle represents the cumulative change in detected lipid classes (increase in DKO mice), a green circle (decrease in DKO mice), or a white circle (no cumulative change). Cumulative changes are computed by counting the number of both increased and decreased levels among the selected discriminant lipid in all lifetime stages. Pathway information was derived from existing literature. Abbreviations: G3P: Glycerol-3-phosphate, PA: Phosphatidic acid, DG: Diacylglycerols, TG: Triacylglycerols, PC: Phosphatidylcholines, PC O-: Ether phosphatidylcholines, PE: Phosphatidylethanolamines, PE O-: Ether phosphatidylethanolamines, LPE: Lysophosphatidylethanolamines, LPC: Lysophosphatidylcholines, PI: Phosphatidyl inositol, HMG CoA: 3-hydroxy-3-methylglutaryl coenzyme A, MUFA: mono-unsaturated fatty acids, PUFA: Poly-unsaturated fatty acids, SM: Sphingomyelin, Cer: Ceramide, HexCer: Hexosylceramide, CK: Choline kinase, ACC: acetyl-CoA carboxylase, ACL: ATP-citrate lyase, FAS: Fatty acid synthase, SCD1: Stearoyl-CoA desaturase-1, UGCG: uridine diphosphate-glucose ceramide glucosyltransferase.

## Materials and Methods

### Experimental Design

*Dicer* ^*flox/flox*^ *Pten* ^*flox/flox*^ *Amhr2* ^*cre/+*^ DKO females and *Dicer* ^*flox/flox*^ *Pten* ^*flox/flox*^ control females that do not carry *Amhr2* ^*cre/+*^ were generated, with the genotypes confirmed by PCR amplification of DNA. Mice were housed in the Baylor College of Medicine vivarium in dedicated mouse rooms in microisolator cages. When animals reached eight weeks of age, serum samples were collected from mice every two weeks until the end of the study or humane endpoint for sacrifice. When a DKO mouse with an advanced-stage cancer was determined to be severely sick, the mouse was anesthetized for the last blood collection via cardiac puncture, and euthanized. The submandibular vein was chosen for the serial blood collection by alternating cheek sides following a valid animal protocol (AN-716). A total of 100-200 µl blood sample was collected into a BD serum separator, allowed for 30 minutes clotting time, and followed by centrifugation and serum collection. Collected serum samples were stored at −80 °C for further metabolomics analysis. DKO mice were sacrificed for this study in accordance to the animal protocol approved by the institutional animal Care and Use Committee (IACUC) at Baylor College of Medicine. Samples from 15 DKO mice (*n* = 231) and 15 control mice (*n* = 238) were used for lipidomics analyses. Prior to data analysis, timepoints for each sample collected were converted into a percentage lifetime metric with the following mathematical formula:

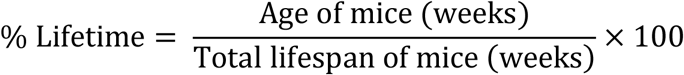

The % lifetimes were then binned into five categories: 0-30% (stage I), 30-45% (stage II), 45-60% (stage III), 60-75% (stage IV), and 75-100% (stage V).

### Reagents

Optima LC-MS grade water, 2-propanol, acetonitrile, formic acid (99.5+%), ammonium formate, and ammonium acetate were purchased from Fisher Chemical (Fisher Scientific International, Inc. Pittsburgh, PA) and used to prepare chromatographic mobile phases and solvents for extraction. Isotopically labeled lipid standards (Table S5) were purchased from Avanti Polar Lipids (Alabaster, AL) and used to prepare the lipid internal standard mixture.

### Sample Preparation

The lipid extraction solvent was prepared by adding 700 µL of the isotopically labeled lipid standard mixture (Table S5) to 42 mL of 2-propanol. Serum samples were thawed on ice, followed by the extraction of non-polar metabolites. The extraction procedure was carried out by adding the prepared extraction solvent to 10-25 µL serum sample in a 3:1 ratio. Following this step, samples were vortex-mixed for 30 s and centrifuged at 13,000 rpm for 7 min. The supernatant was transferred to LC vials and stored at −80 °C until analysis, which was performed within a week. A blank sample, prepared with LC-MS grade water, underwent the same sample preparation process as the serum samples. A pooled quality control (QC) sample was prepared by adding 2-5 µL aliquot of supernatant to each serum sample. This QC sample was analyzed every 10 injections to assess LC-MS instrument stability through the course of the experiment. Samples were run in a randomized order on consecutive days.

### UHPLC-MS Analysis

Reverse-phase (RP) ultra-high performance liquid chromatography-mass spectrometry (UHPLC-MS) analysis was performed with a Thermo Accucore C30, 150 × 2.1 mm, 2.6 µm particle size column mounted in a Vanquish LC coupled to an Orbitrap ID-X Tribrid mass spectrometer (ThermoFisher Scientific). The mobile phases and chromatographic gradients used are described in Supplementary Table S6. MS data were acquired in positive and negative ion modes in the 150-2000 *m/z* range with a 120,000 mass resolution setting. The most relevant MS parameters are provided in the supplementary section Table S7. Samples were kept at 4 °C in the autosampler during LC-MS analysis while the column temperature was set to 50 °C. An injection volume of 2 µL was used for all runs. For lipid annotation, MS/MS experiments were performed using the Thermo Scientific AcquireX data acquisition workflow. Tandem MS data were acquired at a resolution of 30,000 and an isolation window of 0.4 *m/z*. Precursor ions were fragmented with HCD and CID activation methods. For HCD, stepped normalized collision energy (NCE) of 15, 30, and 45 and a CID collision energy of 40 were used to fragment the precursor ions.

### UHPLC-MS Data Processing

Spectral features (described as *m/z*, retention time pairs) were extracted with Compound Discoverer v3.2 (ThermoFisher Scientific) from the raw files. This procedure included retention time alignment of chromatographic peaks, peak picking, peak area integration, and compound area correction using a QC-based regression curve. The sample blank injection was used to remove background peaks: features with less than five times the peak area of corresponding features in the sample blank were marked as background signals and removed from the dataset. Additionally, features that were not present in at least 50% of the QC sample injections or had a relative standard deviation (RSD) of more than 30% in the QC injections were removed from the dataset.

### Lipid Annotation

Lipid annotation was conducted for selected spectral features detected following filtering. The exact masses and MS/MS spectra of all features were first matched against a curated in-house lipid spectral database. For features of interest that did not have matches in the local database, the generated elemental formulas, exact masses, and MS/MS spectra were matched against databases such as Lipid Maps (*64*) and mzCloud (*65*). A total of 1070 species, which included fatty acids, glycerophospholipids, sphingolipids, and glycerolipids, were successfully annotated with this approach and used for further analysis. The complete dataset of annotated species is available through the Metabolomics Workbench, as described above.

### Global Lipidome Analysis

To investigate alterations at the lipidome level, fold changes were computed by taking the base two logarithmic ratio of the lipid abundances for DKO mice to the DKO control mice 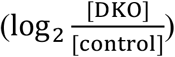. Statistically significant lipids were identified *via* Welch’s *T*-test (DKO *n*=221, DKO control *n*=238) followed by a Benjamini-Hochberg correction using the Statsmodel library (v. 0.12.2). Eighty-seven lipids with *q* < 0.05 were identified as significant. These lipids features were log-transformed (log_2_*X*) and auto-scaled prior to unsupervised machine learning. Principal component analysis (PCA), kernel PCA (kPCA), and t-distributed stochastic neighbor embedding (t-SNE) were performed with the sci-kit learn library (v. 0.24.1). In addition, uniform manifold approximation and projection (UMAP) were performed using the umap library (v. 0.5.1). A two-step pipeline was set up to identify the best hyperparameters for kPCA. First, a kPCA dimensionality reduction to the first two components, followed by a logistic regression classifier, then GridSearchCV in the sci-kit learn library was used to select the best kernel and gamma value for the algorithm. The gamma value selected was 0.03, while the kernel used was the radial basis function (RBF). For t-SNE, the following hyperparameters were used: perplexity= 4, early exaggeration=10. Perplexity controls how the balance between the local and global structure of the data, while early exaggeration is the factor that increases the attractive forces between data points. Time-resolved lipid changes were computed by comparing the five lifetime stages of DKO and DKO control mice with a Welch’s *T*-test. Lipids with *p* < 0.05 were identified as significant. In addition, overlapping significant features in the time-resolved univariate test were identified using an upset plot library (v. 0.6.0). Significant lipids that appeared in at least three lifetime stages were screened as potential prognostic circulating lipids for ovarian cancer.

### Lipidome Longitudinal Analysis

Fold changes, as described above, were computed for 87 lipids with *q* < 0.05, and hierarchical clustering analysis (HCA) was then used to identify clusters of lipidomic trajectories using those fold changes. Each row of the dataset is equivalent to the fold change values over the five lifetime stages for a given lipid feature. The goal of this analysis was to cluster lipids that have a similar trend over time. HCA was performed using the SciPy library (v. 1.6.2). The distance hyperparameter, that is the distance between two observations (lipids), used was the correlation metric, which is defined as follows:

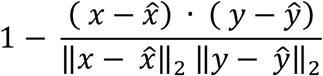

Where *x* and *y* are two lipid features.

The second hyperparameter, the linkage hyperparameter, is the measure of the distance between two clusters to be merged. Complete linkage was used – this method computes the maximum distance between any single data point in the first cluster and any single data point in the second cluster, which is defined as follows:

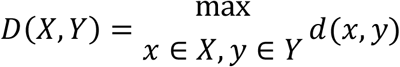

The algorithm then fuses clusters that have the shortest distance between each other. Where *d*(*x,y*) is the distance between lipids *x*∈*X* and *y*∈*Y* and *X* and *Y* are two sets of lipid clusters. Four lipid clusters were identified to have biologically meaningful trends over time. The longitudinal lipid changes of the four lipid clusters were visualized using the Holoview Python library (v. 1.14.6). The correlation network graphs of the four clusters were plotted using Plotly (v. 5.3.1) and networkX (v. 2.5). Lipids with *r* ≥0.5(Pearson’s correlation coefficient) are displayed with a link on the network graphs.

### Machine Learning Classification Methods

#### Feature selection

For each lifetime stage, only lipid features with *P* values < 0.05 (Welch’s *T*-test) were retained. Furthermore, one feature was retained for every two highly correlated lipid features (Pearson’s correlation, *r* > 0.8). Samples were divided into a training set (70% of total samples) and a test set (30% of total samples). Lipid features were selected by fitting the training datasets with a meta-transformer for selecting features based on importance weights. In this case, random forests were used, and features were ranked *via* their Gini index feature importance score. The features with a Gini index greater or equal to the mean of all Gini indices were the final lipid features selected for classification purposes. The number of trees used for the random forest classifiers was a hundred, and all samples were autoscaled prior to feature selection with random forests. Feature selection was carried out with the SelectFromModel function and Random Forest classifier in the sci-kit learn library (v. 0.24.1).

#### ML algorithms

Classification tasks were performed by training machine learning models to discriminate DKO from DKO control mice using the features selected as described above. The machine learning algorithms used included logistic regression, random forest, *k*-nearest neighbors, support vector machines, and a voting ensemble classifier. The default parameters of Python’s sci-kit learn machine learning library (v. 0.24.1) were used. As indicated above, 70% of samples were used for training purposes, with a 5-fold cross-validation method, while the remaining 30% were used as the test set. The classifiers were evaluated using the area under the curve of the receiver operating characteristic curve (AUC ROC) metric. ROC is a probability curve that plots the true positive rate (TPR) against the false positive rate (FPR) at various threshold values. This feature makes it an unbiased metric score, particularly for an unbalanced dataset.

#### Logistic regression

Logistic regression is a regression algorithm used for classification purposes, in this case, binary classification (DKO *vs*. DKO control mice). It is an extension of linear regression, as it computes a weighted sum of input features in addition to a bias term. However, instead of outputting a numeric value as in linear regression, the numeric value is passed through a sigmoid function that computes a probability 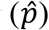 value between 0 and 1.

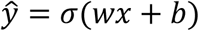

Where *σ*(∙) is the logistic function, ***w*** is the weights/vector coefficient, *x* is lipid features,***b*** is the bias term, and *ŷ* is the final prediction. ***w*** and ***b*** are the parameters set during training and are used to classify samples of the test sets. Probability values are stratified as described below:

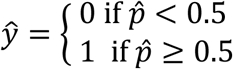

In our case, samples with 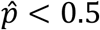 were classified as control animals, while 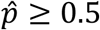 were classified as DKO animals.

#### Random forest classification

Random forests are an ensemble of decision trees. A decision tree takes the form of an inverted tree, starting with a root node at the top, with the node split by lipid features into internal nodes, culminating with the leaf node. While lipid features split each node, as indicated, the leaf nodes give the final classification of either DKO or DKO control mice. Decision trees are assembled to form the random forest via bootstrap aggregation, which reduces prediction variance by random sampling of training samples with replacement. The algorithm also introduces additional randomness during tree construction by using a random subset of features to search for the best features to split the node, resulting in greater tree diversity. For this work, the number of trees in the forest is a hundred, and the quality of node split is measured by the Gini impurity.

#### Support vector machines

The goal of support vector machines (SVM) is to identify a separating hyperplane ***b*** + ***w***^***T***^***x*** that will discriminate two classes of samples with the widest possible margins. Where ***w*** is the weights or coefficient vector, ***b*** is the bias term, and ***x*** is the feature value. This goal is accomplished by learning the ***w*** and ***b*** terms during training with the following equation:

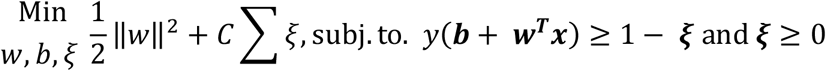

Where *C* is a regularization parameter that penalizes or accommodates *ξ, ξ* is the slack variable that allows for a soft margin classification, allowing some training data to fall within the SVM margin. Therefore, the goal is to minimize the weights, bias, and slack variables, subject to a correct prediction while accommodating the slack variables. In this work, *C* was set to 1. A kernelized SVM was used to transform datasets that are not linearly separable to a higher-dimensional space, where they may be linearly separable. The kernel used in this work is the radial basis function kernel which is defined below:

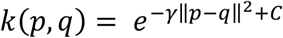

Where *p* and *q* represent data points and *γ* is the kernel coefficient. After training, given a test sample *x*, its prediction score can be obtained with OV score = *b* + *wx*. If the ovarian cancer (OV) score ≤ 0, the sample is classified as control mice, and vice-versa.

#### k-Nearest Neighbors (k-NN)

k-NN is a non-parametric supervised learning algorithm using an instance-based learning method. It simply stores training data instances and computes votes based on the majority class of the *k* nearest neighbors. The number of neighbors selected was five in this work, and a uniform weight function was used. That is, all points in each neighborhood were weighted equally.

#### Voting classifier

Because we selected machine learning models with different inductive biases, we explored an ensemble method voting classifier. The estimators for the voting classifier include all the ML models prior described: logistic regression, random forests, SVM, and k-NN. In addition, soft voting was performed, using average predicted probabilities to predict class labels.

### Prognostic Lipid Discovery Methods and Survival Analysis

Feature selection was performed by a lifetime stage-resolved volcano plot analysis. This involves plotting the -log_10_*P*-value (Welch’s *T*-test, DKO lifetime stages II-V *vs*. DKO stage I) against the log_2_FC (Fold change, DKO lifetime stage II-V *vs*. DKO stage I). Lipid features with *P*-values < 0.05 and at least one log_2_FC for each comparison pair were identified as significant. Volcano plot analysis was performed using the Bioinfokit library (v. 2.0.8). Overlapping significant features in the DKO volcano plot analysis were identified using an upset plot via the Upset python library (v. 0.6.0). Lipids that were significant in at least three of the four DKO lifetime stages comparisons were screened as potential prognostic circulating lipids for ovarian cancer. In addition, significant lipids in at least three lifetime stages comparison of DKO *vs*. control lifetime stages comparisons were also screened.

The selected lipids were used to split the DKO samples into two groups using the median split method. For the last serum collection before mice death or end of the study, the DKO samples with less than or equal to the median of the lipid’s relative abundance were designated as the “low metabolite level” group. In contrast, the DKO samples with greater than the median of the lipid’s relative abundance were designated the “high metabolite level” group. Furthermore, the survival function *S*(*t*) = *P*(*T* > *t*), which is the probability that a mouse survives longer than some specified time *t*, was computed using the Kaplan Meier (KM) estimate described in the equation below:

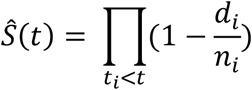

Where *d*_*i*_ is the number of mice death events at time *t*, while *n*_*i*_ is the number of mice at risk of death prior to time *t*. The log-rank test (*p* < 0.05) was used to determine if the differences between KM curves were statistically significant. In addition, the restricted mean survival time (RMST) is defined below:

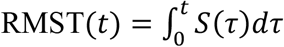

This metric was used to compare two survival curves by measuring the area under the survival curve, which is a measure of “time lost.” Kaplan Meier estimates and the RMST was also used to compute and compare the survival curves of DKO *vs*. control mice, respectively. Finally, the hazard curves were computed using the Nelson-Aalen estimate, and all survival analysis methods in this work were performed using the Python lifelines library (v. 0.26.3).

### Statistical Analysis

Computational analysis was carried out as indicated in the respective sections above using the Python 3.8.8 programming language. NumPy (v. 1.20.1) was used for numerical computations, the Pandas (v. 1.2.4) library was used to perform data handling, and data manipulation, Matplotlib (v. 3.3.4), Plotly (v. 5.3.1) and Holoview (v. 1.14.6) were used for data plotting and visualization.

## Author contributions

Conceptualization: OOB, FMF, DAG

Methodology: OOB, FMF, SS, DAG

Investigation: SS, OOB, SGM

Software: OOB,

Visualization: OOB,

Supervision: FMF, DAG

Writing—original draft: OOB, FMF

Writing—review & editing: OOB, FMF, SS, JK, DAG, SGM, MP

Funding acquisition: FMF

## Data and materials availability

Data generated in this study are available through the NIH Metabolomics Workbench (http://www.metabolomicsworkbench.org/) with project ID PR001457 (study ID ST002276 [http://dx.doi.org/10.21228/M8D133]). Code is available on GitHub: https://github.com/obifarin/DKO-lipidomics

## Supplementary Materials

**Figure S1.**
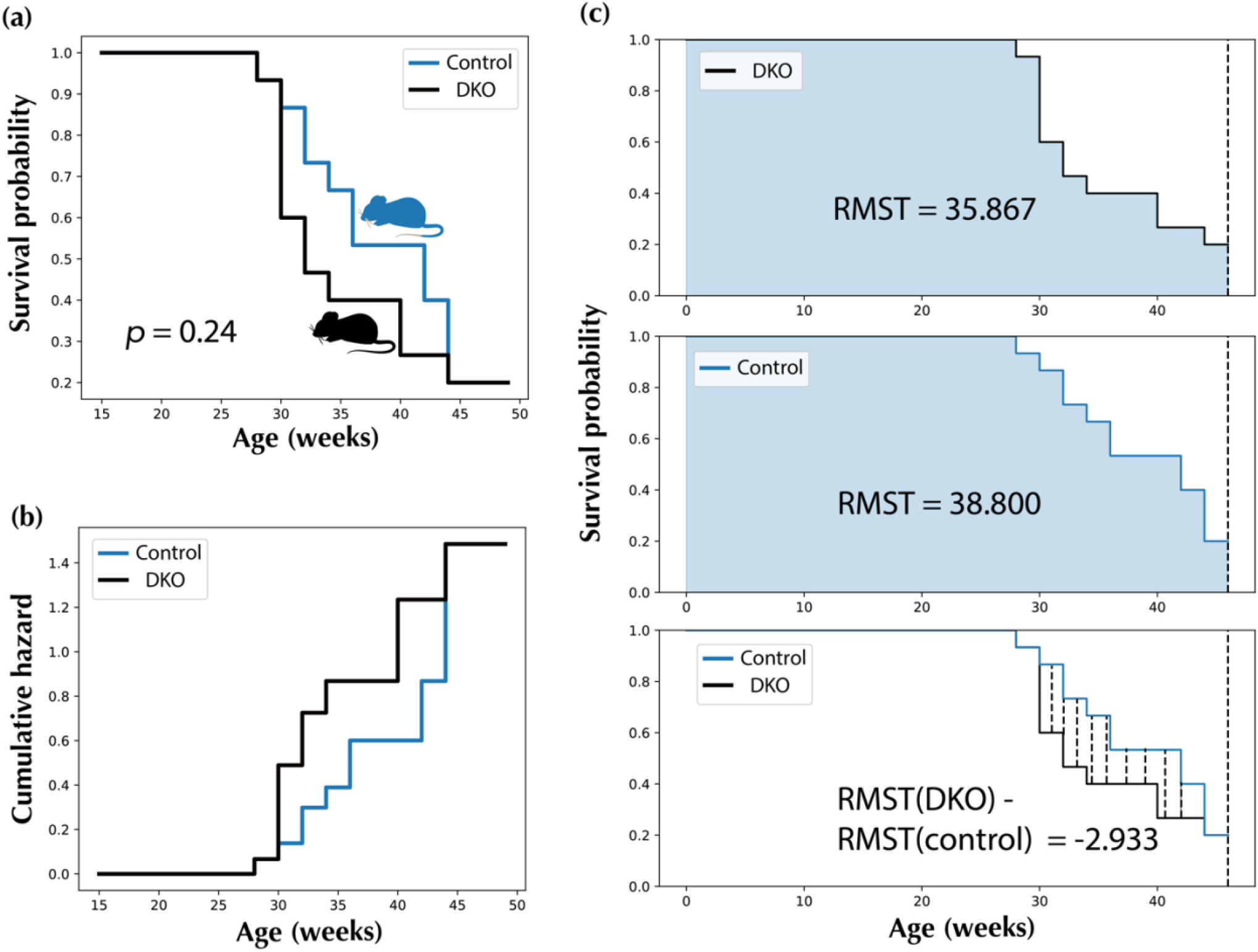
Survival analysis comparison of DKO and DKO control mice. (a) Kaplan-Meier survival curve estimate, DKO *vs*. DKO control mice. (b), Nelson-Aalen hazard curve estimate, DKO *vs*. DKO control mice. (c), Restricted Mean Survival Times (RMST), DKO *vs*. DKO control mice.

**Figure S2.**
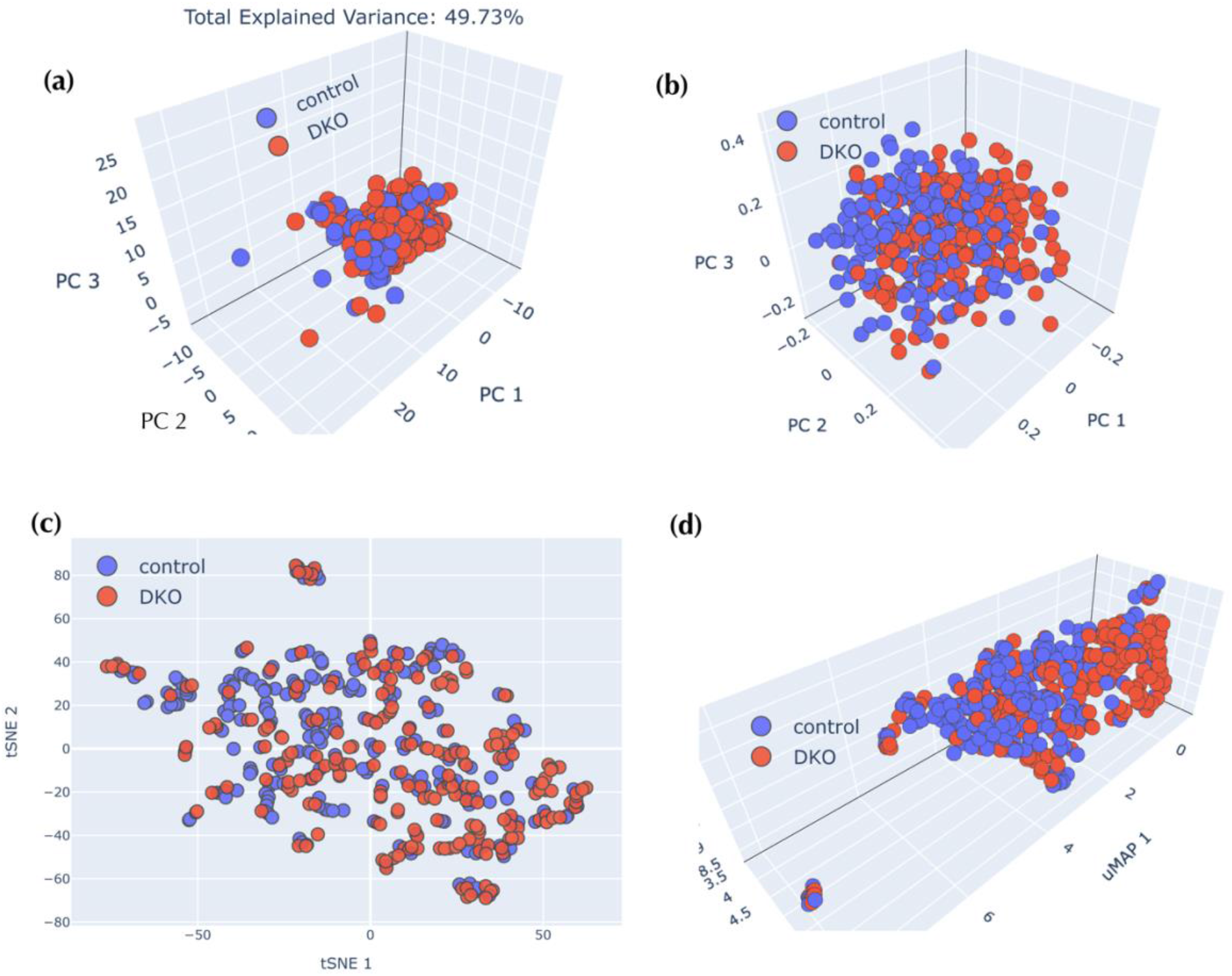
DKO and DKO control mice comparison *via* unsupervised learning methods. (a), PCA score plot. (b), Kernel PCA score plot. (c), tSNE score plot. (d), UMAP score plot. Eighty-seven statistically significant lipid abundances were used for unsupervised learning, and all-time points were combined.

**Figure S3.**
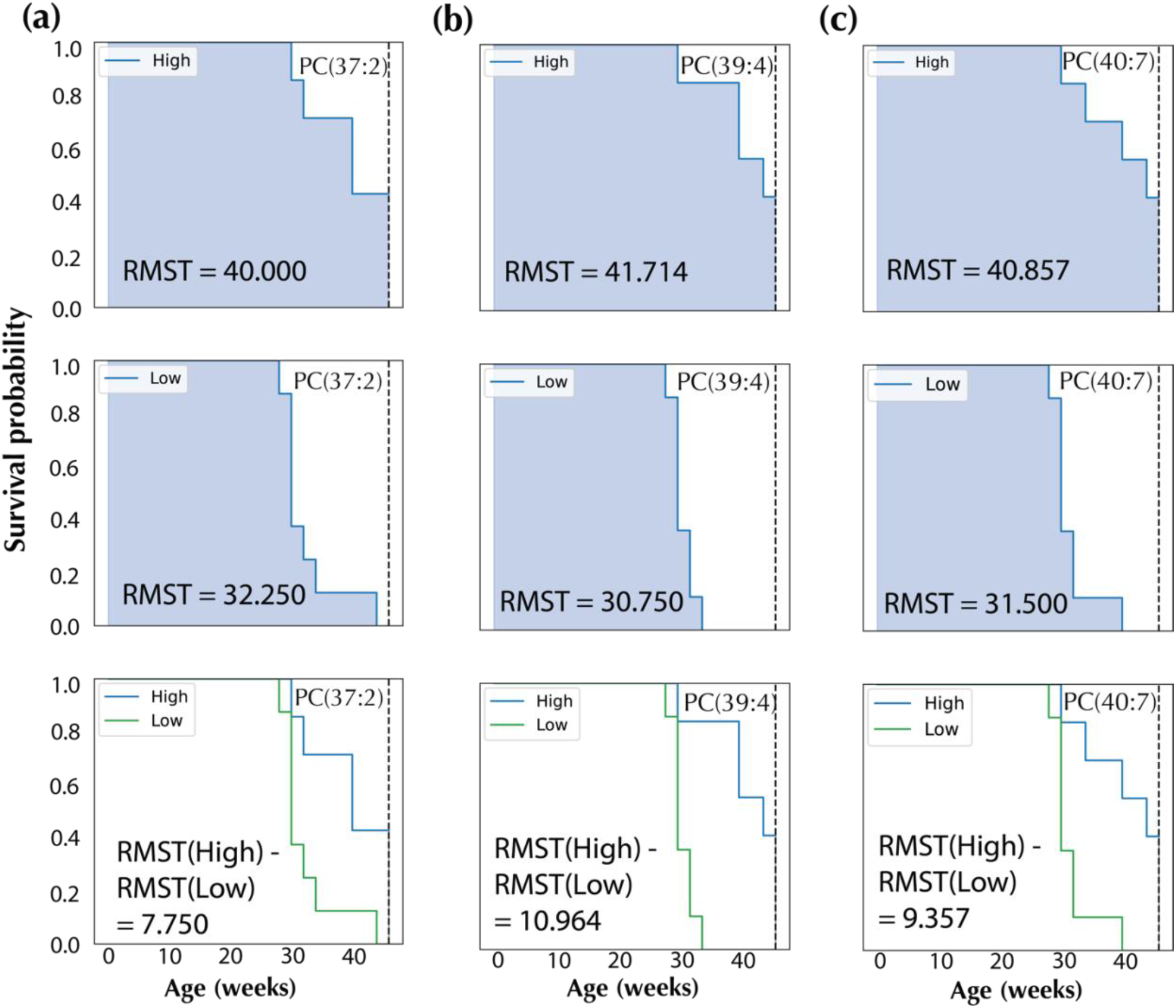
Restricted Mean Survival Times (RMST) plots for all prognostic lipid candidates. (a), PC(37:2). (b), PC(39:4). (c), PC(40:7).

**Table S1.** Eighty-seven statistically significant (*q* < 0.05) lipids for the DKO *vs*. DKO control comparison, all time points combined. DG: Diacylglycerols, TG: Triacylglycerols, FA: Fatty acids, HexCer: Hexosylceramides, LPC: Lysophosphatidylcholines, LPE: Lysophosphatidylethanolamines, PC: Phosphatidylcholines, PC-O: Ether phosphatidylcholines, PE: Phosphatidylethanolamines, PE-O: Ether phosphatidylethanolamines, PI: Phosphatidylinositols, Cer: Ceramides, and SM: Sphingomyelins.

| ID | Retention Time [min] | Lipids | Adduct |
| --- | --- | --- | --- |
| 24 | 1.54 | FA(14:1) | [M-H]<sup>-</sup> |
| 1472 | 4.67 | PE(O-34:3) | [M-H]<sup>-</sup> |
| 1111 | 4.69 | Cer(d33:1) | [M+CH<sub>3</sub>COOH-H]<sup>-</sup> |
| 966 | 4.70 | Cer(d34:1) | [M-H]<sup>-</sup> |
| 1149 | 4.72 | Cer(d34:1) | [M+CH<sub>3</sub>COOH-H]<sup>-</sup> |
| 1217 | 6.83 | Cer(d40:1) | [M-H]<sup>-</sup> |
| 1454 | 6.64 | Cer(d41:2) | [M+CH<sub>3</sub>COOH-H]<sup>-</sup> |
| 1297 | 6.79 | Cer(d42:2) | [M-H]<sup>-</sup> |
| 1290 | 6.46 | Cer(d42:3) | [M-H]<sup>-</sup> |
| 1504 | 6.46 | Cer(d42:3) | [M+CH<sub>3</sub>COOH-H]<sup>-</sup> |
| 111 | 2.37 | FA(18:1) | [M-H]<sup>-</sup> |
| 1473 | 4.33 | HexCer(d34:1) | [M-H]<sup>-</sup> |
| 1761 | 4.09 | HexCer(d34:1) | [M+CH<sub>3</sub>COOH-H]<sup>-</sup> |
| 1762 | 4.32 | HexCer(d34:1) | [M+CH<sub>3</sub>COOH-H]<sup>-</sup> |
| 2078 | 6.83 | HexCer(d42:1) | [M+CH<sub>3</sub>COOH-H]<sup>-</sup> |
| 2532 | 6.83 | HexCer(d42:1) | [M-H]<sup>-</sup> |
| 2065 | 6.41 | HexCer(d42:2) | [M-H]<sup>-</sup> |
| 2522 | 6.41 | HexCer(d42:2) | [M+CH<sub>3</sub>COOH-H]<sup>-</sup> |
| 4260 | 6.51 | HexCer(d42:2) | [M-H]<sup>-</sup> |
| 2415 | 6.41 | HexCer(d42:2) | [M+H<sub>2</sub>CO<sub>2</sub>-H]<sup>-</sup> |
| 257 | 2.03 | FA(20:4-2OH) | [M-H]<sup>-</sup> |
| 52 | 2.02 | FA(16:1) | [M-H]<sup>-</sup> |
| 1870 | 4.26 | PC(16:0_16:0) | [M+H<sub>2</sub>CO<sub>2</sub>-H]<sup>-</sup> |
| 2050 | 5.45 | PC(16:0_18:0) | [M+H<sub>2</sub>CO<sub>2</sub>-H]<sup>-</sup> |
| 2110 | 3.76 | PC(16:0_18:2) | [M+CH<sub>3</sub>COOH-H]<sup>-</sup> |
| 1679 | 4.14 | PC(16:0_18:2) | [M+H<sub>2</sub>CO<sub>2</sub>-H]<sup>-</sup> |
| 1751 | 4.51 | PC(16:0_18:2) | [M+H<sub>2</sub>CO<sub>2</sub>-H]<sup>-</sup> |
| 2265 | 5.90 | PC(18:0_18:0) | [M+H<sub>2</sub>CO<sub>2</sub>-H]<sup>-</sup> |
| 2163 | 3.56 | PC(16:0_20:5) | [M+H<sub>2</sub>CO<sub>2</sub>-H]<sup>-</sup> |
| 2225 | 5.32 | PC(17:0_18:2) | [M+CH<sub>3</sub>COOH-H]<sup>-</sup> |
| 4126 | 4.56 | PC(18:0_20:4) | [2M+H<sub>2</sub>CO<sub>2</sub>-H]<sup>-</sup> |
| 2725 | 5.09 | PC(18:0_22:4) | [M+CH<sub>3</sub>COOH-H]<sup>-</sup> |
| 2626 | 5.20 | PC(20:0_20:4) | [M+H<sub>2</sub>CO<sub>2</sub>-H]<sup>-</sup> |
| 1876 | 5.18 | PC(O-16:0_16:0) | [M+CH<sub>3</sub>COOH-H]<sup>-</sup> |
| 1849 | 2.05 | PC(O-17:1_15:1) and PC(O-16:1_18:1) | [M+CH<sub>3</sub>COOH-H]<sup>-</sup> |
| 2038 | 2.37 | PC(O-16:0_18:1) | [M+CH<sub>3</sub>COOH-H]<sup>-</sup> |
| 2205 | 4.51 | PC(O-18:1_18:2) | [M+CH<sub>3</sub>COOH-H]<sup>-</sup> |
| 2187 | 4.50 | PC(O-16:1_20:3) | [M+CH<sub>3</sub>COOH-H]<sup>-</sup> |
| 2432 | 5.33 | PC(O-18:0_20:3) | [M+CH<sub>3</sub>COOH-H]<sup>-</sup> |
| 2411 | 4.52 | PC(O-18:1_20:3) | [M+CH<sub>3</sub>COOH-H]<sup>-</sup> |
| 2412 | 5.04 | PC(O-18:1_20:3) | [M+CH<sub>3</sub>COOH-H]<sup>-</sup> |
| 2400 | 4.52 | PC(O-18:1_20:4) and PC(O-16:0_22:5) | [M+CH<sub>3</sub>COOH-H]<sup>-</sup> |
| 2587 | 4.33 | PC(O-18:1_22:6) and PC(O-22:7_18:0) | [M+CH<sub>3</sub>COOH-H]<sup>-</sup> |
| 1650 | 4.36 | (16:0_20:4) | [M-H]<sup>-</sup> |
| 1651 | 4.17 | (16:0_20:4) | [M-H]<sup>-</sup> |
| 1623 | 4.35 | PE(O-15:1_22:5) | [M-H]<sup>-</sup> |
| 1699 | 4.33 | PE(O-18:3_20:4) | [M-H]<sup>-</sup> |
| 1641 | 4.38 | PE(O-22:8_18:0) and PE(O-18:2_22:6) | [M-H]<sup>-</sup> |
| 1837 | 4.36 | PE(O-22:8_18:0) and PE(O-18:2_22:6) | [M-H]<sup>-</sup> |
| 2631 | 3.50 | PI(18:1_20:4) | [M-H]<sup>-</sup> |
| 349 | 2.38 | Prostaglandin A1 ethyl ester | [M-H]<sup>-</sup> |
| 1912 | 4.00 | SM(d36:2) | [M+CH<sub>3</sub>COOH-H]<sup>-</sup> |
| 2439 | 6.14 | SM(d41:2) | [M+CH<sub>3</sub>COOH-H]<sup>-</sup> |
| 2557 | 7.17 | SM(d42:1) | [M+CH<sub>3</sub>COOH-H]<sup>-</sup> |
| 78 | 2.21 | FA(17:1) | [M-H]<sup>-</sup> |
| 886 | 2.27 | LPE(20:0) | [M-H]<sup>-</sup> |
| 5443 | 9.32 | DG(40:0) | [M+NH<sub>4</sub>]<sup>+</sup> |
| 10366 | 7.56 | TG(60:12) | [M+NH<sub>4</sub>]<sup>+</sup> |
| 5438 | 9.29 | Campesterol Ester(18:2) | [M+NH<sub>4</sub>]<sup>+</sup> |
| 5439 | 9.35 | Campesterol Ester(18:2) | [M+NH<sub>4</sub>]<sup>+</sup> |
| 5344 | 6.87 | Cer(d18:1_24:1) | [M+H]<sup>+</sup> |
| 6813 | 6.54 | HexCer(d18:1_24:1) | [M+H]<sup>+</sup> |
| 6573 | 4.80 | PC(37:3) | [M+H]<sup>+</sup> |
| 6839 | 4.95 | PC(38:3) | [M+H]<sup>+</sup> |
| 7105 | 5.26 | PC(39:4) | [M+H]<sup>+</sup> |
| 7357 | 4.52 | PC(40:5) | [M+H]<sup>+</sup> |
| 7604 | 5.05 | PC(41:6) | [M+H]<sup>+</sup> |
| 7565 | 5.26 | PC(41:7) | [M+H]<sup>+</sup> |
| 7815 | 4.08 | PC(42:8) | [M+H]<sup>+</sup> |
| 5618 | 5.22 | PC(O-32:0) | [M+H]<sup>+</sup> |
| 5604 | 4.49 | PC(O-32:1) | [M+H]<sup>+</sup> |
| 6538 | 5.08 | PC(O-38:4) | [M+H]<sup>+</sup> |
| 6539 | 5.09 | PC(O-38:4) | [M+H]<sup>+</sup> |
| 6493 | 4.57 | PC(O-38:5) | [M+H]<sup>+</sup> |
| 7022 | 5.26 | PC(O-40:6) | [M+H]<sup>+</sup> |
| 7023 | 4.63 | PC(O-40:6) | [M+H]<sup>+</sup> |
| 6983 | 4.06 | PC(O-40:7) | [M+H]<sup>+</sup> |
| 6984 | 4.40 | PC(O-40:7) | $[M+H]^+$ |
| 9401 | 7.59 | TG(56:9) | $[M+NH_4]^+$ |
| 8964 | 7.46 | TG(58:11) | $[2M+K]^+$ |
| 9614 | 7.46 | TG(58:11) | $[2M+NH_4]^+$ |
| 10101 | 7.46 | TG(58:11) | $[M+K]^+$ |
| 10226 | 7.65 | TG(58:9) | $[M+NH_4]^+$ |
| 10227 | 7.69 | TG(58:9) | $[M+NH_4]^+$ |
| 10228 | 7.79 | TG(58:9) | $[M+NH_4]^+$ |
| 10230 | 7.99 | TG(58:9) | $[M+NH_4]^+$ |
| 4512 | 9.48 | cholesterol derivative | $[M+NH_4]^+$ |

**Table S2.** Statistically significant lipid features for the comparison between DKO and DKO control mice that were present in at least three lifetime stages. FA: Fatty acids, PC: Phosphatidylcholines, PC-O: Ether phosphatidylcholines, Cer: Ceramides, and SM: Sphingomyelins.

| ID | Retention Time [min] | Lipids |
| --- | --- | --- |
| 201 | 1.13 | 15-deoxy-D-12,14-Prostaglandin A2 |
| 1111 | 4.69 | Cer(d33:1) |
| 1454 | 6.64 | Cer(d41:2) |
| 2163 | 3.56 | PC(16:0_20:5) |
| 2409 | 4.77 | PC(18:0_20:4) |
| 2725 | 5.09 | PC(18:0/22:4) |
| 1849 | 2.05 | PC(O-17:1_15:1) and PC(O-16:1_18:1) |
| 2587 | 4.33 | PC(O-18:1_22:6) and PC(O-22:7_18:0) |
| 349 | 2.38 | Prostaglandin A1 ethyl ester |
| 2238 | 6.32 | SM(t39:0) or SM(d39:0-OH) |
| 4431 | 1.64 | FA(18:3) |
| 5618 | 5.22 | PC(O-32:0) |
| 5604 | 4.49 | PC(O-32:1) |
| 6538 | 5.08 | PC(O-38:4) |
| 6539 | 5.09 | PC(O-38:4) |
| 6493 | 4.57 | PC(O-38:5) |
| 7022 | 5.26 | PC(O-40:6) |
| 7023 | 4.63 | PC(O-40:6) |
| 6983 | 4.06 | PC(O-40:7) |

**Table S3.**
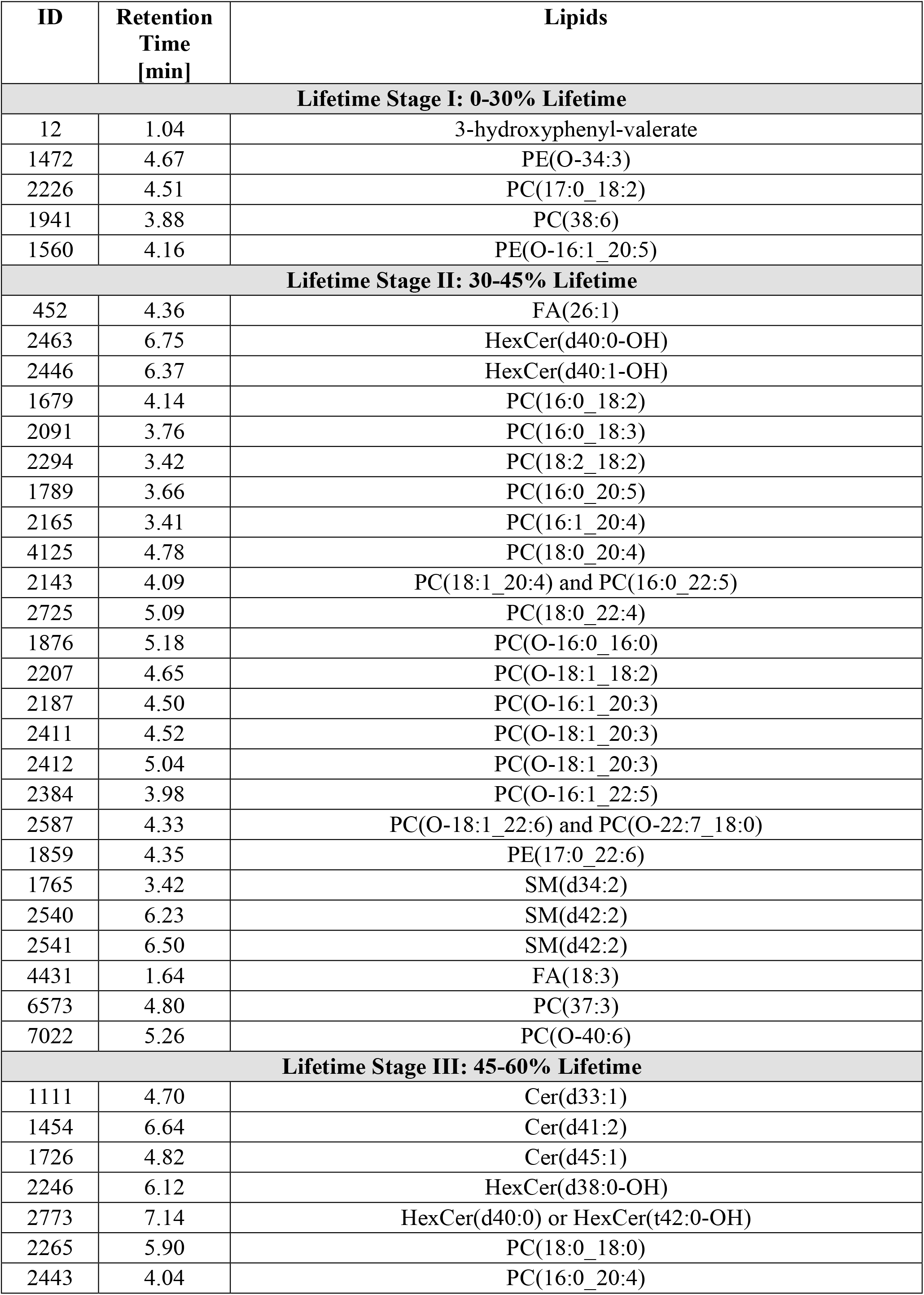
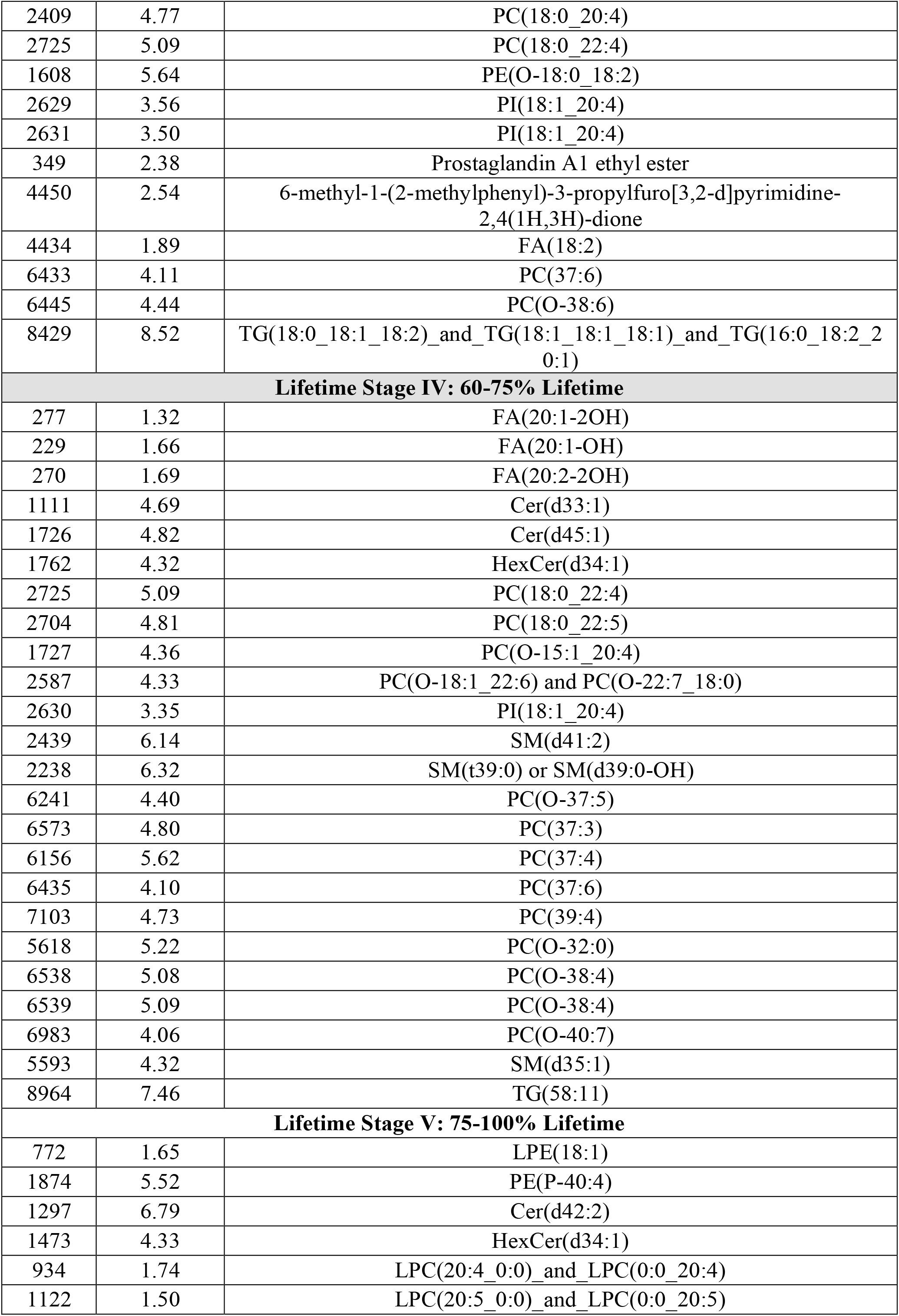
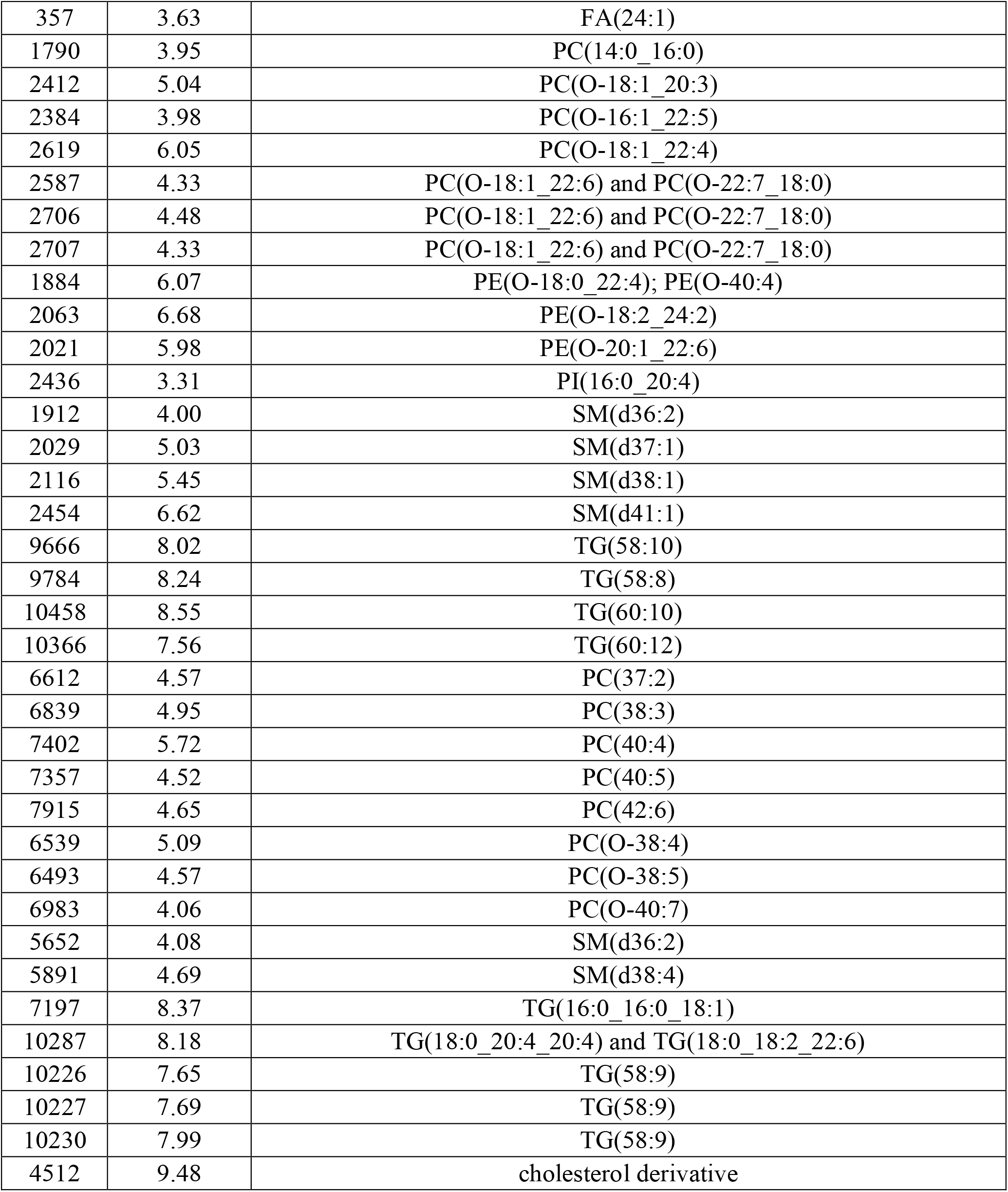
Lipids selected *via* machine learning for each percentage lifetime stage.

**Table S4.** Machine learning results for DKO classification. k-NN: k-Nearest Neighbors, RF: Random Forests, SVM: Support Vector Machine, Voting: Voting Classifier. CV: cross-validation. All scores are ROC AUC.

| Machine learning algorithm | Training set CV scores | Test set scores |
| --- | --- | --- |
| **Lifetime Stage I: 0-30% Lifetime** |  |  |
| Logistic regression | 0.78(±0.16) | 0.74 |
| RF | 0.82(±0.17) | 0.80 |
| k-NN | 0.77(±0.24) | 0.80 |
| SVM | 0.73(±0.19) | 0.74 |
| Voting | 0.76(±0.20) | 0.80 |
| **Lifetime Stage II: 30-45% Lifetime** |  |  |
| Logistic regression | 0.76(±0.21) | 0.66 |
| RF | 0.87(±0.09) | 0.70 |
| k-NN | 0.79(±0.12) | 0.66 |
| SVM | 0.80(±0.11) | 0.62 |
| Voting | 0.82(±0.13) | 0.66 |
| **Lifetime Stage III: 45-60% Lifetime** |  |  |
| Logistic regression | 0.66(±0.08) | 0.85 |
| RF | 0.76(±0.13) | 0.77 |
| k-NN | 0.81(±0.09) | 0.80 |
| SVM | 0.80(±0.06) | 0.78 |
| Voting | 0.77(±0.09) | 0.82 |
| **Lifetime Stage IV: 60-75% Lifetime** |  |  |
| Logistic regression | 0.80(±0.08) | 0.47 |
| RF | 0.83(±0.13) | 0.66 |
| k-NN | 0.76(±0.14) | 0.54 |
| SVM | 0.78(±0.07) | 0.57 |
| Voting | 0.82(±0.11) | 0.54 |
| **Lifetime Stage V: 75-100% Lifetime** |  |  |
| Logistic regression | 0.90(±0.06) | 0.69 |
| RF | 0.90(±0.04) | 0.63 |
| k-NN | 0.90(±0.06) | 0.67 |
| SVM | 0.90(±0.05) | 0.75 |
| Voting | 0.91(±0.04) | 0.74 |

**Table S5.**
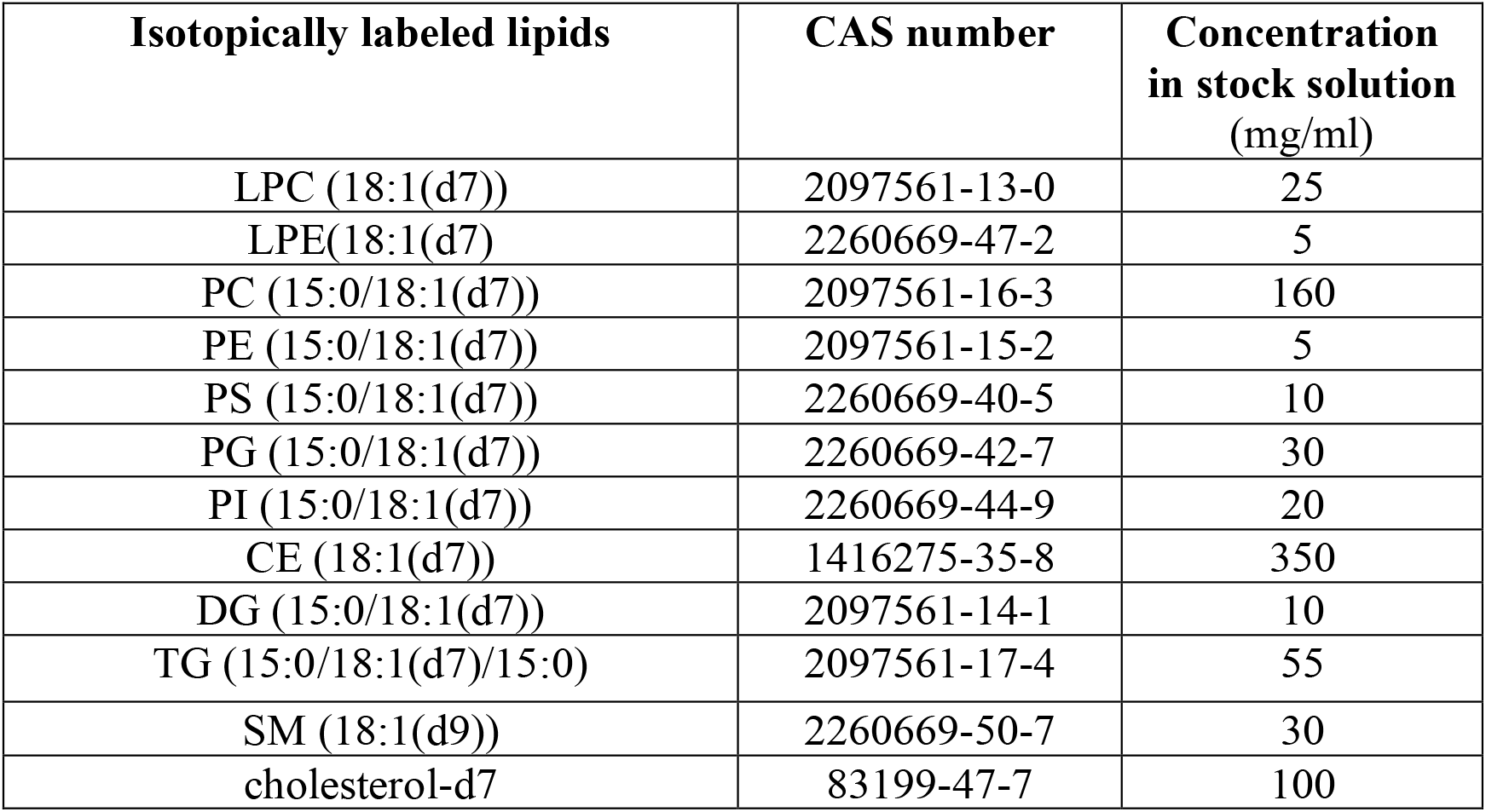
Composition of stable isotope-labeled chemical standards mixture used in UHPLC-MS.

**Table S6.** Chromatographic gradient for RP UHPLC-MS method. For negative ion mode, mobile phase A was 10 mM ammonium acetate with water/acetonitrile (40:60 v/v) and mobile phase B was 10 mM ammonium acetate with 2-isopropanol/acetonitrile (90:10 v/v). For positive ion mode, mobile phase A was 10 mM ammonium formate with water/acetonitrile (40:60 v/v) and 0.1% formic acid. Mobile phase B was 10 mM ammonium formate with 2-isopropanol/acetonitrile (90:10 v/v) and 0.1% formic acid.

| RP UHPLC Gradient |  |  |  |
| --- | --- | --- | --- |
| Time<br>(min) | Mobile<br>phase A | Mobile phase B | Flow rate<br>(ml min<sup>-1</sup>) |
| 0.0 | 80% | 20% | 0.4 |
| 0.0 | 80% | 20% | 0.4 |
| 1.0 | 40% | 60% | 0.4 |
| 5.0 | 30% | 70% | 0.4 |
| 5.5 | 15% | 85% | 0.4 |
| 8.0 | 10% | 90% | 0.4 |
| 8.2 | 0% | 100% | 0.4 |
| 10.5 | 0% | 100% | 0.4 |
| 10.7 | 80% | 20% | 0.4 |
| 12.0 | 80% | 20% | 0.4 |

**Table S7.** MS parameters used for RP UHPLC-MS. Arb: Arbitrary units.

| **MS parameters for RP UHPLC-MS** |  |  |
| --- | --- | --- |
| **MS parameter** | **Positive mode** | **Negative mode** |
| Capillary temperature | 275 °C | 275 °C |
| Spray voltage | + 3.5kV | - 2.5kV |
| Sheath gas flow rate | 40 Arb. | 40 Arb. |
| Auxiliary gas flow rate | 8 Arb. | 8 Arb. |
| Sweep gas flow rates | 1 Arb. | 1 Arb. |
| Vaporizer temperature | 320 °C | 320 °C |

## References

1. A. N. Giaquinto, R. R. Broaddus, A. Jemal, R. L. Siegel, The Changing Landscape of Gynecologic Cancer Mortality in the United States. Obstet Gynecol 139, 440–442 (2022).

2. M. Zou, Y. Du, R. Liu, Z. Zheng, J. Xu, Nanocarrier-delivered small interfering RNA for chemoresistant ovarian cancer therapy. Wiley Interdiscip Rev RNA 12, e1648 (2021).

3. L. A. Torre, B. Trabert, C. E. DeSantis, K. D. Miller, G. Samimi, C. D. Runowicz, M. M. Gaudet, A. Jemal, R. L. Siegel, Ovarian cancer statistics, 2018. CA Cancer J Clin 68, 284–296 (2018).

4. E. Surveillance, and End Results (SEER) Program (http://www.seer.cancer.gov) SEER*Stat Database. (National Cancer Institute, DCCPS, Surveillance Research Program, 2022).

5. M. A. Lisio, L. Fu, A. Goyeneche, Z. H. Gao, C. Telleria, High-Grade Serous Ovarian Cancer: Basic Sciences, Clinical and Therapeutic Standpoints. Int J Mol Sci 20, (2019).

6. J. Kim, E. Y. Park, O. Kim, J. M. Schilder, D. M. Coffey, C. H. Cho, R. C. Bast, Jr., Cell Origins of High-Grade Serous Ovarian Cancer. Cancers (Basel) 10, (2018).

7. J. Kim, D. M. Coffey, C. J. Creighton, Z. Yu, S. M. Hawkins, M. M. Matzuk, High-grade serous ovarian cancer arises from fallopian tube in a mouse model. Proc Natl Acad Sci U S A 109, 3921–3926 (2012).

8. O. Kim, E. Y. Park, D. L. Klinkebiel, S. D. Pack, Y. H. Shin, Z. Abdullaev, R. E. Emerson, D. M. Coffey, S. Y. Kwon, C. J. Creighton, S. Kwon, E. C. Chang, T. Chiang, N. Yatsenko, J. Chien, D. J. Cheon, Y. Yang-Hartwich, H. Nakshatri, K. P. Nephew, R. R. Behringer, F. M. Fernandez, C. H. Cho, B. Vanderhyden, R. Drapkin, R. C. Bast, Jr., K. D. Miller, A. R. Karpf, J. Kim, In vivo modeling of metastatic human high-grade serous ovarian cancer in mice. PLoS Genet 16, e1008808 (2020).

9. J. R. Cantor, D. M. Sabatini, Cancer cell metabolism: one hallmark, many faces. Cancer Discov 2, 881–898 (2012).

10. J. K. Nicholson, J. C. Lindon, Systems biology: Metabonomics. Nature 455, 1054–1056 (2008).

11. K. Bingol, Recent Advances in Targeted and Untargeted Metabolomics by NMR and MS/NMR Methods. High Throughput 7, (2018).

12. M. I. Jordan, T. M. Mitchell, Machine learning: Trends, perspectives, and prospects. Science 349, 255–260 (2015).

13. O. O. Bifarin, D. A. Gaul, S. Sah, R. S. Arnold, K. Ogan, V. A. Master, D. L. Roberts, S. H. Bergquist, J. A. Petros, F. M. Fernandez, A. S. Edison, Machine Learning-Enabled Renal Cell Carcinoma Status Prediction Using Multiplatform Urine-Based Metabolomics. J Proteome Res 20, 3629–3641 (2021).

14. D. A. Gaul, R. Mezencev, T. Q. Long, C. M. Jones, B. B. Benigno, A. Gray, F. M. Fernandez, J. F. McDonald, Highly-accurate metabolomic detection of early-stage ovarian cancer. Sci Rep 5, 16351 (2015).

15. E. I. Braicu, S. Darb-Esfahani, W. D. Schmitt, K. M. Koistinen, L. Heiskanen, P. Poho, J. Budczies, M. Kuhberg, M. Dietel, C. Frezza, C. Denkert, J. Sehouli, M. Hilvo, High-grade ovarian serous carcinoma patients exhibit profound alterations in lipid metabolism. Oncotarget 8, 102912–102922 (2017).

16. H. S. Ahn, J. Yeom, J. Yu, Y. I. Kwon, J. H. Kim, K. Kim, Convergence of Plasma Metabolomics and Proteomics Analysis to Discover Signatures of High-Grade Serous Ovarian Cancer. Cancers (Basel) 12, (2020).

17. S. Plewa, A. Horala, P. Derezinski, E. Nowak-Markwitz, J. Matysiak, Z. J. Kokot, Wide spectrum targeted metabolomics identifies potential ovarian cancer biomarkers. Life Sci 222, 235–244 (2019).

18. X. Wang, X. Zhao, J. Zhao, T. Yang, F. Zhang, L. Liu, Serum metabolite signatures of epithelial ovarian cancer based on targeted metabolomics. Clin Chim Acta 518, 59–69 (2021).

19. C. M. Jones, M. E. Monge, J. Kim, M. M. Matzuk, F. M. Fernandez, Metabolomic serum profiling detects early-stage high-grade serous ovarian cancer in a mouse model. J Proteome Res 14, 917–927 (2015).

20. H. Lusk, J. E. Burdette, L. M. Sanchez, Models for measuring metabolic chemical changes in the metastasis of high grade serous ovarian cancer: fallopian tube, ovary, and omentum. Mol Omics 17, 819–832 (2021).

21. R. S. Freedman, M. Deavers, J. Liu, E. Wang, Peritoneal inflammation - A microenvironment for Epithelial Ovarian Cancer (EOC). J Transl Med 2, 23 (2004).

22. J. Cai, H. Tang, L. Xu, X. Wang, C. Yang, S. Ruan, J. Guo, S. Hu, Z. Wang, Fibroblasts in omentum activated by tumor cells promote ovarian cancer growth, adhesion and invasiveness. Carcinogenesis 33, 20–29 (2012).

23. S. Sah, X. Ma, A. Botros, D. A. Gaul, S. R. Yun, E. Y. Park, O. Kim, S. G. Moore, J. Kim, F. M. Fernandez, Space- and Time-Resolved Metabolomics of a High-Grade Serous Ovarian Cancer Mouse Model. Cancers (Basel) 14, (2022).

24. J. N. van der Veen, J. P. Kennelly, S. Wan, J. E. Vance, D. E. Vance, R. L. Jacobs, The critical role of phosphatidylcholine and phosphatidylethanolamine metabolism in health and disease. Biochim Biophys Acta Biomembr 1859, 1558–1572 (2017).

25. C. Stoica, A. K. Ferreira, K. Hannan, M. Bakovic, Bilayer Forming Phospholipids as Targets for Cancer Therapy. Int J Mol Sci 23, (2022).

26. E. Iorio, A. Ricci, M. Bagnoli, M. E. Pisanu, G. Castellano, M. Di Vito, E. Venturini, K. Glunde, Z. M. Bhujwalla, D. Mezzanzanica, S. Canevari, F. Podo, Activation of phosphatidylcholine cycle enzymes in human epithelial ovarian cancer cells. Cancer Res 70, 2126–2135 (2010).

27. F. Gibellini, T. K. Smith, The Kennedy pathway--De novo synthesis of phosphatidylethanolamine and phosphatidylcholine. IUBMB Life 62, 414–428 (2010).

28. R. J. Niemi, E. I. Braicu, H. Kulbe, K. M. Koistinen, J. Sehouli, U. Puistola, J. U. Maenpaa, M. Hilvo, Ovarian tumours of different histologic type and clinical stage induce similar changes in lipid metabolism. Br J Cancer 119, 847–854 (2018).

29. V. B. O’Donnell, New appreciation for an old pathway: the Lands Cycle moves into new arenas in health and disease. Biochem Soc Trans 50, 1–11 (2022).

30. S. H. Law, M. L. Chan, G. K. Marathe, F. Parveen, C. H. Chen, L. Y. Ke, An Updated Review of Lysophosphatidylcholine Metabolism in Human Diseases. Int J Mol Sci 20, (2019).

31. C. Ke, Y. Hou, H. Zhang, L. Fan, T. Ge, B. Guo, F. Zhang, K. Yang, J. Wang, G. Lou, K. Li, Large-scale profiling of metabolic dysregulation in ovarian cancer. Int J Cancer 136, 516–526 (2015).

32. F. Mazet, M. J. Tindall, J. M. Gibbins, M. J. Fry, A model of the PI cycle reveals the regulating roles of lipid-binding proteins and pitfalls of using mosaic biological data. Sci Rep 10, 13244 (2020).

33. W. Zhao, Y. Qiu, D. Kong, Class I phosphatidylinositol 3-kinase inhibitors for cancer therapy. Acta Pharm Sin B 7, 27–37 (2017).

34. L. Shayesteh, Y. Lu, W. L. Kuo, R. Baldocchi, T. Godfrey, C. Collins, D. Pinkel, B. Powell, G. B. Mills, J. W. Gray, PIK3CA is implicated as an oncogene in ovarian cancer. Nat Genet 21, 99–102 (1999).

35. A. Klippel, M. A. Escobedo, M. S. Wachowicz, G. Apell, T. W. Brown, M. A. Giedlin, W. M. Kavanaugh, L. T. Williams, Activation of phosphatidylinositol 3-kinase is sufficient for cell cycle entry and promotes cellular changes characteristic of oncogenic transformation. Mol Cell Biol 18, 5699–5711 (1998).

36. E. U. Frevert, B. B. Kahn, Differential effects of constitutively active phosphatidylinositol 3-kinase on glucose transport, glycogen synthase activity, and DNA synthesis in 3T3-L1 adipocytes. Mol Cell Biol 17, 190–198 (1997).

37. H. W. Chang, M. Aoki, D. Fruman, K. R. Auger, A. Bellacosa, P. N. Tsichlis, L. C. Cantley, T. M. Roberts, P. K. Vogt, Transformation of chicken cells by the gene encoding the catalytic subunit of PI 3-kinase. Science 276, 1848–1850 (1997).

38. M. W. Holliday, Jr., S. B. Cox, M. H. Kang, B. J. Maurer, C22:0- and C24:0-dihydroceramides confer mixed cytotoxicity in T-cell acute lymphoblastic leukemia cell lines. PLoS One 8, e74768 (2013).

39. G. Grammatikos, N. Schoell, N. Ferreiros, D. Bon, E. Herrmann, H. Farnik, V. Koberle, Piiper, S. Zeuzem, B. Kronenberger, O. Waidmann, J. Pfeilschifter, Serum sphingolipidomic analyses reveal an upregulation of C16-ceramide and sphingosine-1-phosphate in hepatocellular carcinoma. Oncotarget 7, 18095–18105 (2016).

40. L. Chen, H. Chen, Y. Li, L. Li, Y. Qiu, J. Ren, Endocannabinoid and ceramide levels are altered in patients with colorectal cancer. Oncol Rep 34, 447–454 (2015).

41. N. Kozar, K. Kruusmaa, M. Bitenc, R. Argamasilla, A. Adsuar, N. Goswami, D. Arko, I. Takac, Metabolomic profiling suggests long chain ceramides and sphingomyelins as a possible diagnostic biomarker of epithelial ovarian cancer. Clin Chim Acta 481, 108–114 (2018).

42. M. Dany, B. Ogretmen, Ceramide induced mitophagy and tumor suppression. Biochim Biophys Acta 1853, 2834–2845 (2015).

43. S. Sridhar, Y. Botbol, F. Macian, A. M. Cuervo, Autophagy and disease: always two sides to a problem. J Pathol 226, 255–273 (2012).

44. W. Jiang, B. Ogretmen, Autophagy paradox and ceramide. Biochim Biophys Acta 1841, 783–792 (2014).

45. Z. Li, L. Zhang, D. Liu, C. Wang, Ceramide glycosylation and related enzymes in cancer signaling and therapy. Biomed Pharmacother 139, 111565 (2021).

46. H. Yoon, S. Lee, Fatty Acid Metabolism in Ovarian Cancer: Therapeutic Implications. Int J Mol Sci 23, (2022).

47. R. J. Deberardinis, N. Sayed, D. Ditsworth, C. B. Thompson, Brick by brick: metabolism and tumor cell growth. Curr Opin Genet Dev 18, 54–61 (2008).

48. G. Hatzivassiliou, F. Zhao, D. E. Bauer, C. Andreadis, A. N. Shaw, D. Dhanak, S. R. Hingorani, D. A. Tuveson, C. B. Thompson, ATP citrate lyase inhibition can suppress tumor cell growth. Cancer Cell 8, 311–321 (2005).

49. K. Brusselmans, E. De Schrijver, G. Verhoeven, J. V. Swinnen, RNA interference-mediated silencing of the acetyl-CoA-carboxylase-alpha gene induces growth inhibition and apoptosis of prostate cancer cells. Cancer Res 65, 6719–6725 (2005).

50. V. Chajes, M. Cambot, K. Moreau, G. M. Lenoir, V. Joulin, Acetyl-CoA carboxylase alpha is essential to breast cancer cell survival. Cancer Res 66, 5287–5294 (2006).

51. Y. Cai, J. Wang, L. Zhang, D. Wu, D. Yu, X. Tian, J. Liu, X. Jiang, Y. Shen, L. Zhang, M. Ren, P. Huang, Expressions of fatty acid synthase and HER2 are correlated with poor prognosis of ovarian cancer. Med Oncol 32, 391 (2015).

52. U. V. Roongta, J. G. Pabalan, X. Wang, R. P. Ryseck, J. Fargnoli, B. J. Henley, W. P. Yang, J. Zhu, M. T. Madireddi, R. M. Lawrence, T. W. Wong, B. A. Rupnow, Cancer cell dependence on unsaturated fatty acids implicates stearoyl-CoA desaturase as a target for cancer therapy. Mol Cancer Res 9, 1551–1561 (2011).

53. K. M. Gharpure, S. Pradeep, M. Sans, R. Rupaimoole, C. Ivan, S. Y. Wu, E. Bayraktar, A. S. Nagaraja, L. S. Mangala, X. Zhang, M. Haemmerle, W. Hu, C. Rodriguez-Aguayo, M. McGuire, C. S. L. Mak, X. Chen, M. A. Tran, A. Villar-Prados, G. A. Pena, R. Kondetimmanahalli, R. Nini, P. Koppula, P. Ram, J. Liu, G. Lopez-Berestein, K. Baggerly, S. E. L A. K. Sood, FABP4 as a key determinant of metastatic potential of ovarian cancer. Nat Commun 9, 2923 (2018).

54. Y. Liang, H. Han, L. Liu, Y. Duan, X. Yang, C. Ma, Y. Zhu, J. Han, X. Li, Y. Chen, CD36 plays a critical role in proliferation, migration and tamoxifen-inhibited growth of ER-positive breast cancer cells. Oncogenesis 7, 98 (2018).

55. A. J. O’Donnell, K. G. Macleod, D. J. Burns, J. F. Smyth, S. P. Langdon, Estrogen receptor-alpha mediates gene expression changes and growth response in ovarian cancer cells exposed to estrogen. Endocr Relat Cancer 12, 851–866 (2005).

56. H. J. Kim, R. K. Kalkhoff, Sex steroid influence on triglyceride metabolism. J Clin Invest 56, 888–896 (1975).

57. W. R. Hazzard, M. J. Spiger, J. D. Bagdade, E. L. Bierman, Studies on the mechanism of increased plasma triglyceride levels induced by oral contraceptives. N Engl J Med 280, 471–474 (1969).

58. T. O’Brien, T. T. Nguyen, Lipids and lipoproteins in women. Mayo Clin Proc 72, 235–244 (1997).

59. H. Ulmer, W. Borena, K. Rapp, J. Klenk, A. Strasak, G. Diem, H. Concin, G. Nagel, Serum triglyceride concentrations and cancer risk in a large cohort study in Austria. Br J Cancer 101, 1202–1206 (2009).

60. Y. Kikuchi, M. Miyauchi, K. Oomori, T. Kita, I. Kizawa, K. Kato, Inhibition of human ovarian cancer cell growth in vitro and in nude mice by prostaglandin D2. Cancer Res 46, 3364–3366 (1986).

61. B. Diez-Dacal, D. Perez-Sala, A-class prostaglandins: early findings and new perspectives for overcoming tumor chemoresistance. Cancer Lett 320, 150–157 (2012).

62. B. Huang, B. L. Song, C. Xu, Cholesterol metabolism in cancer: mechanisms and therapeutic opportunities. Nat Metab 2, 132–141 (2020).

63. K. J. Helzlsouer, A. J. Alberg, E. P. Norkus, J. S. Morris, S. C. Hoffman, G. W. Comstock, Prospective study of serum micronutrients and ovarian cancer. J Natl Cancer Inst 88, 32–37 (1996).

64. E. Fahy, M. Sud, D. Cotter, S. Subramaniam, LIPID MAPS online tools for lipid research. Nucleic Acids Res 35, W606–612 (2007).

65. R. Mistrik, mzCLOUD: A spectral tree library for the Identification of “unknown unknowns”. Abstracts of Papers of the 255th American Chemical Society National Meeting, (2018).

